# Mobile genetic element insertions drive antibiotic resistance across pathogens

**DOI:** 10.1101/527788

**Authors:** Matthew G. Durrant, Michelle M. Li, Ben Siranosian, Ami S. Bhatt

## Abstract

Mobile genetic elements contribute to bacterial adaptation and evolution; however, detecting these elements in a high-throughput and unbiased manner remains challenging. Here, we demonstrate a *de novo* approach to identify mobile elements from short-read sequencing data. The method identifies the precise site of mobile element insertion and infers the identity of the inserted sequence. This is an improvement over previous methods that either rely on curated databases of known mobile elements or rely on ‘split-read’ alignments that assume the inserted element exists within the reference genome. We apply our approach to 12,419 sequenced isolates of nine prevalent bacterial pathogens, and we identify hundreds of known and novel mobile genetic elements, including many candidate insertion sequences. We find that the mobile element repertoire and insertion rate vary considerably across species, and that many of the identified mobile elements are biased toward certain target sequences, several of them being highly specific. Mobile element insertion hotspots often cluster near genes involved in mechanisms of antibiotic resistance, and such insertions are associated with antibiotic resistance in laboratory experiments and clinical isolates. Finally, we demonstrate that mutagenesis caused by these mobile elements contributes to antibiotic resistance in a genome-wide association study of mobile element insertions in pathogenic *Escherichia coli*. In summary, by applying a *de novo* approach to precisely identify mobile genetic elements and their insertion sites, we thoroughly characterize the mobile element repertoire and insertion spectrum of nine pathogenic bacterial species and find that mobile element insertions play a significant role in the evolution of clinically relevant phenotypes, such as antibiotic resistance.

## Main

Genomic variation is critical for the adaptation of pathogenic bacteria. Successful human pathogens, such as *Escherichia coli* and *Staphylococcus aureus*, are able to acquire adaptive phenotypes, such as antibiotic resistance, through the acquisition of single nucleotide polymorphisms, small insertions and deletions, inversions, duplications, and also through the movement of mobile genetic elements (MGEs). Prokaryotic MGEs come in many forms such as insertion sequence (IS) elements, transposons, integrons, plasmids, and bacteriophages (Stokes and Gillings 2011; Rankin, Rocha, and Brown 2011). Some of these elements can mobilize and insert themselves in a site-specific or random manner throughout the host genome. They are of great interest to the scientific and medical communities, in large part because they can transfer between microbes via horizontal gene transfer, spreading their genetic potential between strains of the same species, and even across species (Juhas 2015; Bloemendaal, Brouwer, and Fluit 2010).

IS elements are among the simplest MGEs, and typically code for only the gene (transposase) necessary for their transposition (Mahillon and Chandler 1998). IS elements can transpose into genes, resulting in insertional mutagenesis and loss of function of the gene (Figure 1f) (Emmanuelle Lerat and Ochman 2004); alternatively, they can transpose into gene regulatory elements and influence expression of neighboring genes (Figure 1f,g). IS elements play a role in antibiotic resistance (Linkevicius, Sandegren, and Andersson 2013), horizontal gene transfer (Ochman, Lawrence, and Groisman 2000), and genome evolution (Plague et al. 2017; Biémont and Vieira 2006; Barrick et al. 2009). In addition to minimal elements, such as insertion sequences, MGEs can contain additional “passenger” genes (Figure 1e). These passenger genes can code for a variety of proteins, including virulence factors, antibiotic resistance genes, detoxifying agents, and enzymes for secondary metabolism (Rankin, Rocha, and Brown 2011).

**Figure 1:**
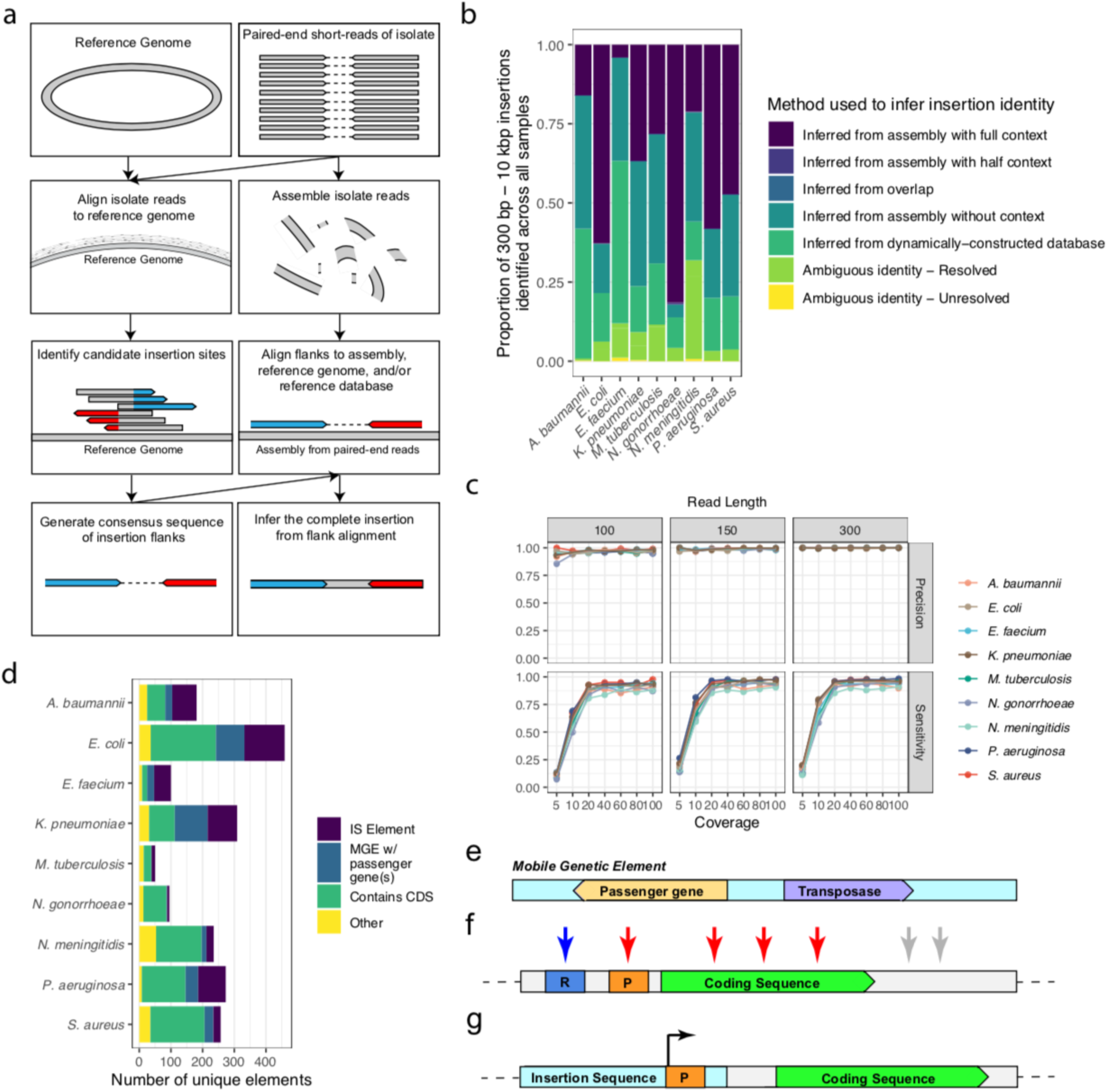
A new approach to identify MGEs from short-read sequencing data. **a**, A schematic representation of the workflow implemented in this study. The analysis requires a reference genome of a given species and short-read sequencing FASTQ files as input. The reads are aligned to the provided reference genome and assembled using third-party software. Candidate MGE insertion sites are identified from the alignment to the reference using our own software suite, called *mustache*. This approach identifies sites where oppositely-oriented clipped read ends are found within 20 bases of each other. Consensus sequences of these flanking ends are then identified, with the assumption being that they represent the flanks of the inserted element. The intervening sequence between the candidate flanks is inferred by aligning flanks to the assembled genome, a reference genome, and our dynamically constructed reference database of all identified MGEs. **b**, The inference method used to characterize the inserted elements of the downloaded data. “Inferred from assembly with full context” indicates that the sequence was found in the expected sequence context within the assembly and is considered one of the highest-confidence inferred sequences. “Inferred from assembly with half context” indicates that the sequence was found in the expected sequence context on one end of the inserted element, and the other element was truncated at the end of a partially assembled contig. “Inferred from overlap” indicates that the sequence was recovered simply by finding an overlap of two paired flanks. Very few elements are recovered by this method, as we are limiting our analysis to only elements greater than 300 base pairs in length. “Inferred from assembly without context” indicates that the sequence identified was recovered from the assembly, but not within the expected sequence context, presumably due to assembly errors. “Inferred from dynamically-constructed database” refers to elements that were not recovered from the assembly but were recovered from the reference genome or from our dynamically constructed database. The database in this case is built from the accumulated inferred sequences found in the assembly and in the reference across all sequenced isolates. “Ambiguous identity - Resolved” indicates that the element was initially is assigned to multiple element clusters but was resolved using several techniques described in the methods. “Ambiguous identity - Unresolved” indicates the the inserted element maps to multiple element clusters and could not be resolved. This excludes these insertions from some of the downstream analyses. **c,** Results of simulations using the *mustache* software tools. We simulated insertions into the nine species analyzed in this study. Reference genomes were mutated with base pair substitutions and short indels at a rate of 0.085 mutations per base pair. An insertion is considered found if both of its flanks are recovered near the expected insertion seite, properly paired with each other, and the consensus flanks show high similarity with the expected inserted sequence. Precision indicates the proportion of reported results that are true simulated MGE insertions, and sensitivity indicates the proportion of simulated MGE insertions that are recovered by *mustache*. **d**, An analysis of the proportion of elements identified in the *mustache* workflow. A “unique element” refers to a unique cluster of elements that are 95% similar across 85% of their respective sequences. The number of elements predicted to be IS elements is shown in purple. “MGE w/ passenger genes” in blue indicates other elements that contain both predicted transposases and one or more passenger genes. “Contains CDS” in green indicates all other elements that contain a predicted CDS, but no transposase. “Other” in yellow indicates all other inserted elements that contain no predicted CDS. **e-g,** Schematic representations of the many ways that MGEs influence biological processes: **e**, through carrying cargo proteins of specific function. **f,** through disrupting functional genomic elements, and **g**, by introducing outwardly-directed promoters, causing up-regulation of adjacent genes. The MGE insertions depicted in (**f**) affect intergenic regulatory elements, such as (R)epressors and (P)romoters, the coding sequence itself, and regions downstream of coding sequences. Blue arrow indicates a probable gain-of-function MGE insertion causing greater expression of the respective gene. Red arrows indicate probable loss-of-function MGE insertion. Grey arrows indicate mutations of unknown function but are likely to be neutral.

Historical estimates suggest that IS movements make up approximately 3% of acquired mutations in adaptive laboratory evolution (ALE) experiments (Conrad, Lewis, and Palsson 2011), but more recent studies have found that the rate of IS mutagenesis can vary depending on growth conditions, such as oxygen exposure (Finn et al. 2017). Under neutral conditions, the rate of IS element insertion in *E. coli* is estimated as 3.5 x 10^−4^ per genome per generation under neutral laboratory conditions (Lee et al. 2016); this IS insertion rate is about 1/3^rd^ the rate of base substitution (Lee et al. 2012). Whereas the majority of studies investigating IS insertion rates have been carried out in *E. coli*, the rate of IS insertions does vary between strains and species (Knöppel et al. 2018; Lee et al. 2016). Functionally, IS elements have been shown to play an important role in antibiotic resistance in studies of small collections of clinical isolates (Olliver et al. 2005; Sun et al. 2016), but a larger survey of IS insertions in clinical isolates has not yet been carried out. Our somewhat narrow understanding of the location and role of IS elements in bacterial adaptation is likely the result of limited awareness of the role of these elements in adaptation, an assumption that short reads cannot accurately identify such insertions, and the lack of flexible and sensitive tools to identify such mutations.

Several approaches exist for identifying IS elements and cargo-carrying MGEs for both prokaryotic and eukaryotic genomes (Barrick et al. 2014; E. Lerat 2010; Xie and Tang 2017; Treepong et al. 2018; Jiang et al. 2015; Adams, Bishop, and Wright 2016; Biswas et al. 2015; Hawkey et al. 2015). For example, a database-dependent approach has been used to identify novel IS elements through performing a BLAST search on completed genome or draft genome assemblies (Siguier et al. 2006), and other approaches relying on split-read alignments can identify MGE insertions with respect to a well-annotated reference genome (Barrick et al. 2014). While these approaches are useful, most are limited due to their dependence on shared homology with known MGEs or genes, or by requiring well-annotated reference genomes using sequenced isolates that closely resemble the reference.

Here, we sought to comprehensively identify complete MGEs, the genes they contained, and their site of insertion with respect to a reference genome from short-read sequencing data. Our approach is flexible, as it can use a database of MGEs when available, but it does not depend entirely on a database of known MGEs and mobile genes. It is sensitive and precise enough to be used on both laboratory samples and environmental isolates. We focus on MGE insertion sites, generate consensus sequences for inserted elements from the clipped ends of locally-aligned reads, infer complete MGEs from sequence assemblies, and build a *de novo* database of elements across all analyzed samples. We combine several sequence inference approaches to identify large insertions, resulting in a highly sensitive and precise overall approach.

By focusing on MGE insertion sites, we answer several questions about these elements that have not been thoroughly addressed in the past. For example, we determine the target-site specificity of each of these MGEs, their level of activity relative to other MGEs within the same species, and the overall MGE insertional capacity of the species of interest. Understanding the overall MGE potential helps us to understand to what extent a bacterial species can adapt by means of MGE activity, either within a single bacterial genome or by way of horizontal gene transfer. Additionally, we can identify genes that are frequently disrupted by MGE insertions within the population, hinting at selective pressures that may be driving these adaptive insertion events

We use our *de novo* approach to analyze 12,419 sequenced isolates of nine pathogenic bacterial species and demonstrate large differences in the overall MGE repertoire and the rate of MGE insertion between species. We characterize MGE insertion hotspots across species and demonstrate that these insertions alter the activity of genes in clinically relevant biological pathways, such as those related to antibiotic resistance. We analyze MGE insertions in the context of adaptive laboratory evolution (ALE) experiments and demonstrate that MGE insertions mediate many of the loss-of-function adaptations that confer resistance to antibiotics. Finally, we use MGE insertions as features in a bacterial genome-wide association study (GWAS) to identify known and potentially novel mechanisms of antibiotic resistance.

### A *de novo* approach to identify and genotype MGE insertions

Several tools exist to identify mobile element insertions from short-read sequencing data of prokaryotic genomes, each with their own strengths and weaknesses (Supplementary Table 1). Most of the available tools have two major limitations: they either require a database of known mobile elements, such as insertion sequences, to initially identify these large insertions (Hawkey et al. 2015), or were built specifically for use in resequencing studies (such as ALE experiments) (Barrick et al. 2014), thus requiring a well-annotated and very closely related reference genome for the organism being studied, an assumption that often fails when studying diverse strains of the same species. We sought to develop a tool that would overcome these limitations and identify large insertions and their genomic position *de novo*, without the need for either a reference database of mobile elements or a closely related reference genome with well-annotated insertion sequences. We applied this tool, which we call *mustache*, in an analysis of thousands of publicly-available sequenced isolates of the prevalent bacterial pathogens *Acinetobacter baumannii, Enterococcus faecium, Escherichia coli, Klebsiella pneumoniae, Mycobacterium tuberculosis, Neisseria gonorrhoeae, Neisseria meningitidis, Pseudomonas aeruginosa*, and *Staphylococcus aureus*. Short-read sequence data for a random subset of these isolates were downloaded from the Sequence Read Archive (SRA) National Center for Biotechnology Information (NCBI) database, and the subsequent analyses summarize the subsets analyzed.

The approach implemented by *mustache* is described schematically in Figure 1a. Briefly, we first align short reads from isolates of a given species to a reference genome of that species using the BWA mem local alignment tool. We found that while the choice of the reference genome used will determine the insertions that are identified, many of the analyses presented in this study are quite robust to reference genome choice (Supplementary Figure 5). Second, we identify candidate insertion sites by searching alignments for reads that have clipped ends at the same site. A read with a clipped end indicates the exact site where the inserted element begins with respect to the reference genome used, and we build a consensus sequence from these clipped ends. Constructing the consensus sequence this way is where our method fundamentally differs from split-read alignment approaches. Third, these high-quality consensus sequences are paired with nearby oppositely-oriented consensus sequences to identify candidate insertions, represented by these partial reconstructions of the inserted element flanks. The genome of each bacterial isolate is then assembled using the SPAdes assembler (Bankevich et al. 2012), candidate insertion site sequences are aligned back to the assembled genome, and the full inserted sequence is inferred from the alignments. Finally, we build a database of such inferred sequences across all analyzed isolates, and we perform a final sequence inference step of all candidate pairs by aligning to this accumulated database, in the event that the sequence could not be inferred in the initial inference step.

Fundamentally, *mustache* overcomes two limitations of prior approaches. First, since repeated sequences such as MGEs impair genome assembly, alignment-based approaches that compare reference and assembled genomes often fail to identify full insertion sequences. As *mustache* does not require that insertion sequences assemble within their expected sequence context, this limitation is overcome. Second, existing split-read approaches typically assume that the MGE of interest exists in the reference genome being used. However, this assumption often fails when analyzing organisms where the introduction of novel genetic material by horizontal gene transfer is common (Alkan, Coe, and Eichler 2011; Barrick et al. 2014). By combining several inference approaches (Figure 1b; Supplementary Figure 1), *mustache* increases the overall sensitivity and confidence in the accuracy of the inferred sequence. By focusing on reconstructing a high-quality consensus sequence of the inserted element from clipped ends, we can use clipped ends as short as one base pair to support the existence of an insertion at a given site.

It is critical that insertion discovery tools are both precise and sensitive. To evaluate the performance characteristics of *mustache*, we estimated its precision and sensitivity using simulations (Figure 1c). We simulated 32 MGE insertions per isolate genome, using 16 species-specific IS elements chosen at random, and 16 randomly generated sequences between 300 and 10,000 base pairs in length. We used DWGSIM to mutate the reference genome with short indels and point mutations at two mutation rates - 0.001 and 0.085 per base pair (Homer, 2017). We find that at 40x coverage and a mutation rate of 0.085, we have at least 83% sensitivity to detect MGE insertions for all species tested, and over 87% sensitivity at 100x coverage. At a mutation rate of 0.001, sensitivity is over 88% across all coverages greater than 40x. A major strength of our approach is that it filters out small insertions and substitutions that can be mistaken as large insertions, resulting in precision that exceeds 98% when coverage is greater than 40x. ANOVA tests of all simulated samples with 40x coverage or greater indicate that species (*F*_*9,2145*_ = 106; *P* < 2 x 10^−16^), mutation rate (*F*_*1,2145*_ = 199, *P* < 2 x 10^−16^), and read length (*F*_*2,2145*_ *=* 44, *P* < 2 x 10^−16^) all influence sensitivity, but that coverage does not (*F*_*3,2145*_ = 2.5; *P* = 0.0614). Precision is also influenced by species (*F*_*9,2145*_ = 2.7; *P* = 0.006), mutation rate (*F*_*1,2145*_ = 241, *P* < 2 x 10^−16^), and read length (*F*_*2,2145*_ *=* 143, *P* < 2 x 10^−16^), but not coverage (*F*_*3,2145*_ = 0.77; *P* = 0.5). Differences between sensitivity and precision of our approach across species are likely due to genomic differences such as higher number of repetitive regions, as we filter out reads mapping ambiguously within each genome. Differences due to read length and mutation rate should be considered when comparing multiple isolates, although these differences are quite small (Figure 1c). We also compared the performance of *mustache* to a recently published tool, panISa (Treepong et al. 2018), and we find no significant difference in sensitivity above 40x coverage (ANOVA test; *F*_*1,4304*_ = 2.5; *P* = 0.11), but *mustache* has improved sensitivity at coverage below 40x (ANOVA test; *F*_*1,3225*_ = 39; *P* =4.9 x 10^−10^; Supplementary Figure 2). Notably, the precision of mustache is much more stable across different levels of coverage compared to *panISa*. Moreover, *mustache* can infer the complete inserted sequence from an assembly or reference genome, whereas *panISa* relies on a web-based BLAST of the ISfinder database to infer the identity of the inserted element.

While *mustache* appears to be highly accurate, it should be noted that we did not simulate all of the many genomic differences that could exist between isolates, such as inversions or more complex combinations of structural changes. For the purpose of this analysis, we limit our analysis to only those insertion sites whose inferred sequence was between 300 bp and 10 kbp and were inserted into the bacterial chromosome. Additionally, we see differences in the confidence of our predictions between species; for example, more than 50% of *E. coli* insertions across all 1,471 isolates were inferred with “high confidence” by this method, as opposed to *E. faecium* where less than 10% could be inferred with “high confidence” by this approach. Despite these limitations, the simulation studies carried out demonstrate that *mustache* is sensitive and precise.

### Characterizing the MGE repertoire of pathogenic bacterial species

MGEs are known to contribute to antibiotic resistance in certain pathogens, such as *E. coli* and *A. baumannii (*Adams et al. 2008; Linkevicius, Sandegren, and Andersson 2013). Of the nine pathogenic species we analyzed in this study, several have developed high levels of antibiotic resistance in recent years. Thus, understanding their MGE repertoire, common sites of insertion, and their contribution to antibiotic resistance is of interest to researchers and clinicians alike. All of the isolates we analyzed were downloaded at random from the SRA database, a publicly-available database provided by the NCBI. They were filtered according to several quality control parameters (see methods), leaving us with a total of 12,419 sequenced isolates in total (Supplementary Table 2).

We ran *mustache* on all sequenced isolates, and we limited our analysis to only MGEs in the size range from 300 bp to 10 kbp that inserted into the bacterial chromosome. We clustered and annotated the genes, including transposases, encoded in these elements (Supplementary Tables 3-5). Overall, we predict that 25.6% of the identified elements from all nine species sequenced are IS elements, with 53% of the *E. faecium* elements being classified as IS elements, compared to 7.3% of *N. gonorrhoeae* elements (Figure 1d). Another 16.5% of all identified elements are predicted to be MGEs with passenger proteins, defined as elements that contain a predicted transposase and one or more other predicted proteins. The remaining 57.9% of elements contain no transposase, and likely represent one-off insertion events, or deletions that are specific to the reference genome used but were not deleted in the isolate.

Next, we analyzed the number of times that we observed a given sequence element inserted at different loci across the genome (Figure 2a). We found that *A. baumannii* and *E. coli* had the highest number of high-activity MGEs, with 46 elements in *A. baumannii* and 45 elements in *E. coli* appearing at more than 10 positions across all analyzed samples. *N. gonorrhoeae* was on the opposite extreme, with no elements being detected at over 10 positions, although this may be partially due to the fact that we only analyzed 895 *N. gonorrhoeae* isolates compared to 1,471 *E. coli* isolates. Nonetheless, these differences across species suggest distinct differences in MGE diversity and overall activity.

**Figure 2:**
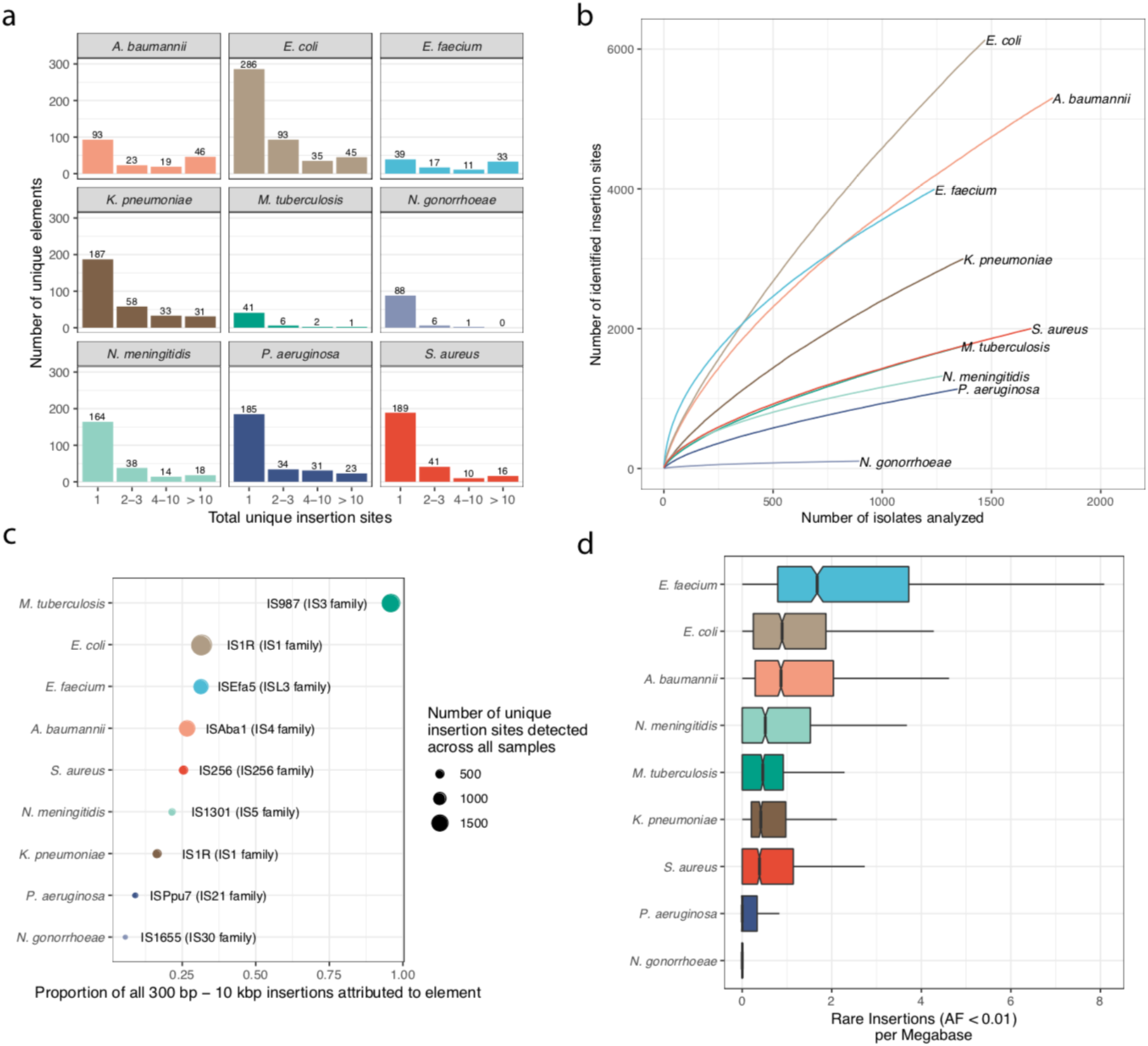
Bacterial species vary considerably in their overall MGE repertoire and rate of insertion. **a**, The number of unique sequence elements identified, binned by species and total unique insertion sites. All 300 bp – 10 kbp sequence elements identified in the workflow were clustered using CD-HIT-EST at 90% similarity across 85% of the sequence. The number of unique insertion sites for each element refers to the number of unique sites where members of a given element cluster can be found across all isolates analyzed for each species. **b**, The number of new insertions identified as additional isolates are analyzed. **c,** An analysis of the most MGEs found in each species, defined by the total number of unique insertion sites where the element is found. The x-axis indicates the proportion of all detected 300 bp - 10kbp insertions that are attributed to the element for each species. The size of each point indicates the total number of unique insertion sites detected for the respective element across all isolates. **d,** Notched boxplots of the number of rare insertions detected across all samples for each species, adjusted by megabase of genome. A rare insertion is defined as one that has an allele frequency (AF) < 0.01 (an insertion that was identified in less than 1% of all samples). The total number of rare insertions identified per isolate was then adjusted by the number of genomic sites in the sequenced isolates that had non-zero coverage, and then multiplied by 1 megabase. Notches indicate 1.58 x IQR / sqrt(n), a rough 95% confidence interval for comparing medians (Mcgill, Tukey, and Larsen 1978).

We find that as more isolates are analyzed, more unique insertions are identified, but that the rate of increase varies from species to species (Figure 2b). *E. coli* is on the high end of the distribution, with 6,131 unique insertions identified after analyzing 1,471 isolates, suggesting that it not only has a high diversity of MGEs but that they are quite active in the population. In contrast, only 106 unique insertions were detected in *N. gonorrhoeae* across all 895 analyzed isolates, indicating that MGE insertions are not a common adaptive strategy for this species. This does not, however, account for differences in genome size across species, which likely partially explains differences in the rate of insertion. To adjust for genome size, we analyzed the number of rare insertions (those with allele frequency < 0.01 in the population) per megabase of covered genome for each isolate (Figure 2d). When adjusting for genome length in this manner, we find that *E. faecium* has the highest number of rare insertions per megabase of reference genome with a mean 2.56 (95% CI, 2.41-2.71) across all analyzed samples, while *N. gonorrhoeae* and *P. aeruginosa* have means of 0.11 (95% CI, 0.06-0.15) and 0.26 (95% CI, 0.23-0.28), respectively. There are definite outliers, such as an *E. faecium* isolate (SRA accession SRR646281) with 70 detected rare insertions (31.8 per covered megabase), and an *E. coli* isolate (SRR3180793) with 93 detected rare insertions (24.7 per covered megabase). Here, we see that the number of MGE insertions continues to increase and that estimates for the rate of insertion vary considerably across species. These estimates can inform researchers going forward as they consider sequencing and analyzing more pathogenic isolates belonging to these species.

We find that the elements responsible for the most insertions are IS elements for each species, as expected (Figure 2c). We find that the IS987 element is responsible for 96% of all detected insertions in *M. tuberculosis*, with 1,673 insertions attributed to this element. The IS1 element is responsible for 32% of all detected insertions in *E. coli*, with 1,938 insertions attributed to it. On the low end we have IS1655 in *N. gonorrhoeae* which accounts for 5.7% (6/106) of detected insertions, and ISPpu7 in *Pseudomonas aeruginosa* which accounts for 9.0% (103/1137) of detected insertions. To our knowledge, these are the first MGE insertional activity estimates for these species, and they can help us to prioritize the overall importance of MGE insertions in each species’ genome evolution. In summary, we have demonstrated that the MGE repertoire and rate of insertion varies significantly across different pathogenic species; however, as a general rule, as more isolates are analyzed, more novel insertions are identified.

### Characterizing MGE passenger proteins

While the most minimal MGE contains only a transposase, which is a gene encoding its own excision and repositioning in the genome, many MGEs contain additional genes referred to as passenger genes. For most species, the majority of the elements appearing in more than one position in the genome contain an open-reading frame (ORF) that codes for a protein homologous to known transposases (Figure 3a). As the number of unique insertions associated with an element increases, the probability that it contains a transposase tends to increase. This strongly indicates that these elements we have identified are in fact mobile. However, other elements do not contain such transposases and appear frequently throughout the genome, such as Group II introns seen in *E. coli, K. pneumoniae, E. faecium*, and *P. aeruginosa* isolates (Fig. 3c). We find that 67 MGEs identified that contain two or more predicted ORFs that could not be annotated using the annotation software Prokka, indicating that their function may be as yet unknown (Figure 3b,d).

**Figure 3:**
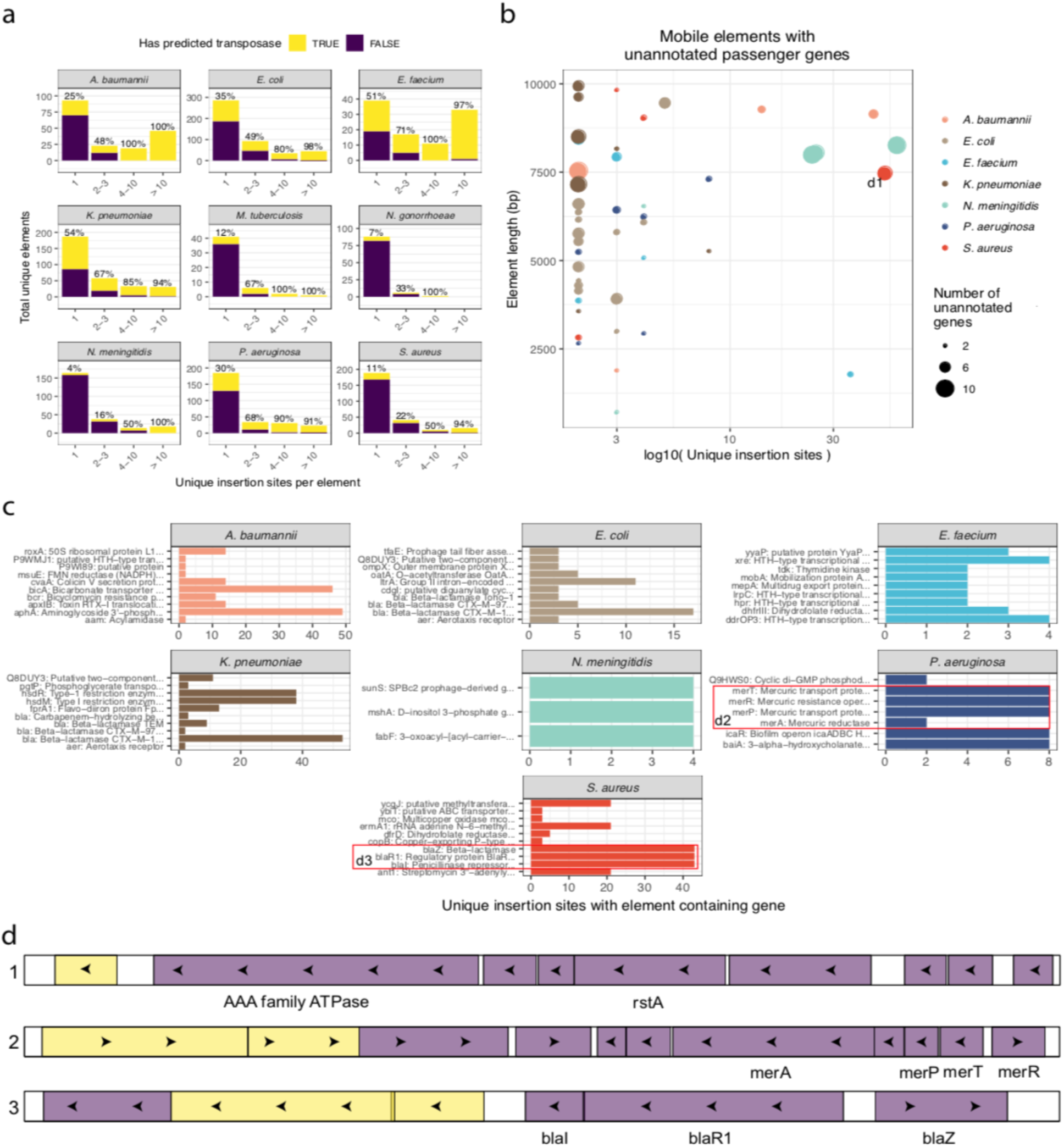
Many identified MGEs include passenger genes, some of which are largely uncharacterized. **a**, The number of immobile (1 unique insertion site) and mobile (> 1 unique insertion site) elements containing a predicted transposase. The percentage over each bar indicates the percentage of elements in each group that contain a predicted transposase. The bins along the x-axis indicate the number of unique insertion sites where elements are found. **b,** Mobile elements with two or more uncharacterized proteins. On the x-axis is the log of the number of unique insertion sites attributed to the element, and on the y-axis is the nucleotide length of the corresponding element. The radius of each point corresponds with the total number of uncharacterized ORFs found in the element (see methods). The labeled point ‘d1’ indicates the first element represented in panel d. **c**, The number of unique insertion sites identified where a mobile element containing each labeled passenger protein was found. Ties in the number of unique sites generally indicates that the passenger proteins are on the same element. The labels “d2” and “d3” in the *P. aeruginosa* and *S. aureus* panels refer to the second and third elements in panel d. **d**, Three examples of mobile elements carrying passenger proteins. The first example corresponds with the *S. aureus* element labelled “1” in b, which contains several uncharacterized ORFs, and includes a AAA family ATPase and transcriptional regulatory protein rstA. The second example is a mercuric resistance mobile element found in *P. aeruginosa*, one of several mercuric resistance elements identified, highlighted in c. The third example is a beta-lactamase containing mobile element found in *S. aureus*, an example of one of several such beta-lactamase-containing mobile elements identified. Yellow rectangles are coding sequences with predicted transposase activity, and purple rectangles are all other predicted coding sequences.

Many of the MGEs that we identified contained passenger ORFs with well-annotated functions (Figure 3c,d). Several elements contain known antibiotic resistance genes as annotated by ResFinder (Zankari et al. 2012), including: 15 *S. aureus* elements coding for aminoglycoside, beta-lactam, macrolide, phenicol, and trimethoprim resistance genes; 10 *E. coli* elements coding for aminoglycoside and beta-lactam resistance genes; 5 *E. faecium* elements coding for aminoglycoside, macrolide, oxazolidinone, phenicol, tetracycline, and trimethoprim resistance genes; one *N. meningitidis* element coding for an aminoglycoside resistance gene, and one *A. baumannii* element coding for an aminoglycoside resistance gene.

Using the annotation software Prokka (Torsten Seemann 2014), we identified 183 elements that contained one or more annotated passenger genes and were found inserted in more than one location, and 73 of these elements also contained at least one predicated transposase. These elements contain many many well-studied mobile genes, such as the resistance genes mentioned above, as well streptomycin 3’-adenylyltransferases, colicin V secretion proteins, bicyclomycin resistance proteins, genes involved in mercuric resistance (*merP, merR, merT, merA*) (Hamlett et al. 1992), and those involved in restriction modification systems (*hsdS, hsdR, hsdM*) (N. E. Murray et al. 1982). A variety of other MGE passenger genes were identified in our workflow, such as bicarbonate transporter protein *bicA*, which appears in a MGE at 46 different sites throughout the *A. baumannii* reference genome across all samples analyzed. In summary, our tool has identified MGEs containing passenger proteins of known and unknown function, shedding light on the phenotypic changes that may accompany these insertion events.

### Analysis of MGE insertion sites reveals functionally significant insertion hotspots

The approach we have taken allows us not only to identify MGEs *de novo*, but it also identifies their sites of insertion with base-pair precision with respect to the reference genome. This allows us to investigate the role that MGEs play in genomic evolution, identifying insertion hotspots that may indicate functionally important genes and pathways. We performed an analysis of MGE insertion hotspots using all unique MGE insertions in each species analyzed (Figure 4a). With the exception of *N. gonorrhoeae*, which had too few insertions to analyze, we identified several MGE insertion hotspots for all species, with 635 being identified in total (Supplementary Table 7). We find that 236 of the hotspots appear to directly overlap with predicted coding sequences, indicating loss-of-function, while 183 are upstream of the nearest gene, and 216 are downstream of the nearest gene, which are more difficult to functionally interpret.

**Figure 4:**
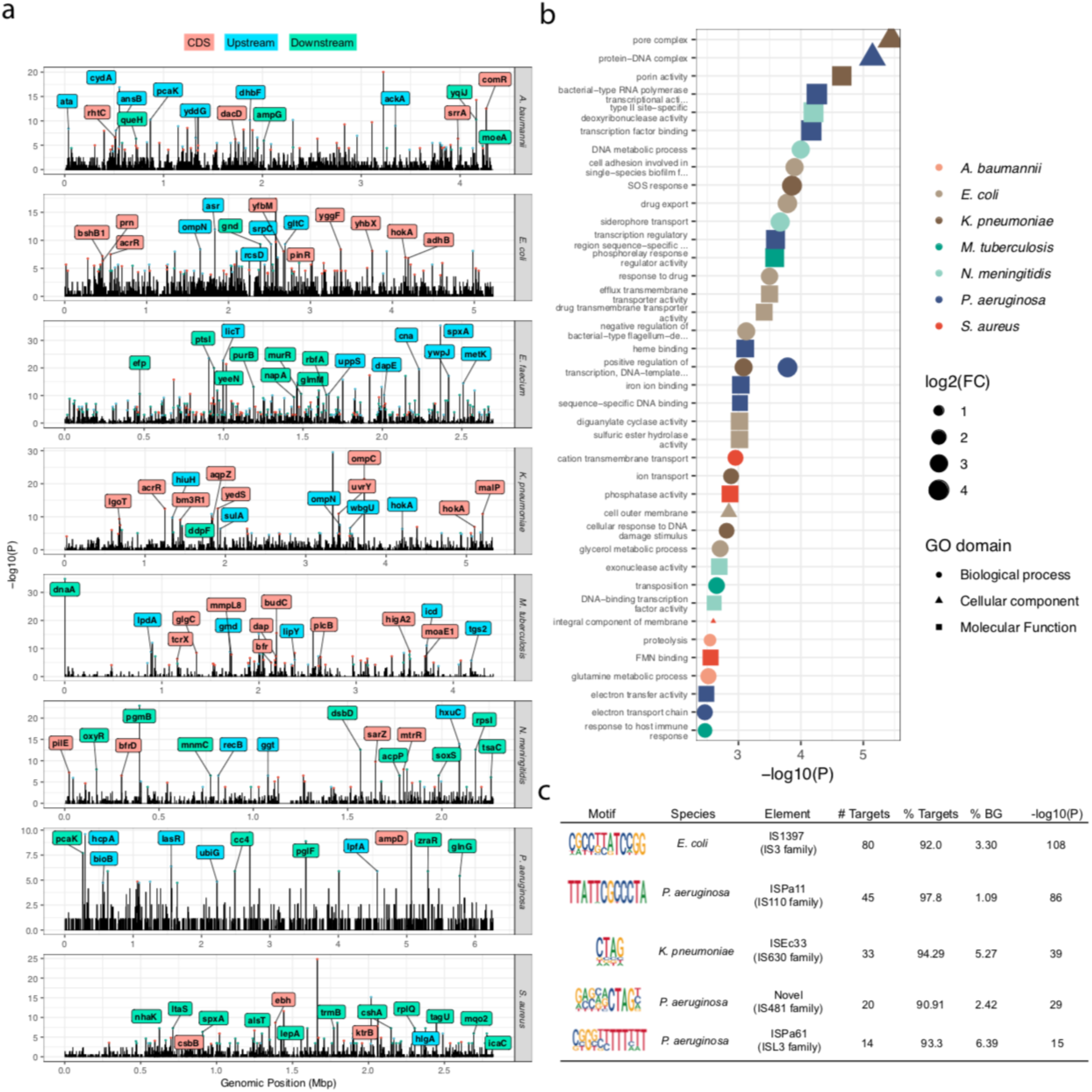
MGE insertion hotspots occur near functionally important genes, and insertion sites reveal target-site specificity. **a**, Analysis of MGE insertion hotspots found on each species’ chromosome. Briefly, 500 bp windows were moved across the genome in 50 bp increments, and peaks are tested using a dynamic Poisson distribution to identify windows statistically enriched for unique MGE insertions. Hotspots were assigned to coding sequences by choosing the coding sequence closest to the center of each hotspot. All hotspots with q-value < 0.05 are indicated with the colored points. The 15 most significant hotspots associated with well-annotated coding sequences are shown in the text labels. Blue labels/points indicate that the hotspot center is upstream of the listed coding sequence, red labels/points indicate that the hotspot center is within the coding sequence itself, and green labels/points indicate that the hotspot center is found downstream of the listed coding sequence. **b**, Gene Ontology (GO) enrichment analysis of predicted coding sequences near MGE hotspots. All coding sequences with q-value < 0.05 were tested for enrichment of each GO term using a hypergeometric test. **c**, Examples of target-sequence identified for five different MGEs with high target-sequence specificity. Motifs identified using the HOMER software. The “Element” column is based on BLAST searches to ISfinder database. “# Targets refers to the number of unique insertion sites analyzed for each MGE to identify motifs. “% Targets” indicates the percentage of target sites containing the motif. “% BG” indicates the percentage of randomly chosen background sequences that contain the motif.

We sought to identify genes that are frequently near MGE insertion hotspots both within and across species. After assigning each hotspot to the closest predicted coding sequence, we clustered all translated proteins by 40% amino acid similarity and 70% sequence coverage. We then identified homologous genes within the same species that were near insertion hotspots, as well as homologous genes that were near hotspots in multiple species. Three such genes were near multiple hotspots both within and across species: *hok/sok* toxin-antitoxin system component, with four homologs near hotspots in *E. coli* and two near hotspots in *K. pneumoniae*; outer membrane porins N and C (*ompN/ompC)* in *E. coli* and *K. pneumoniae*, with *ompN* near a hotspot in *E. coli*, and both *ompC* and *ompN* being near hotspots in *K. pneumoniae*; and Regulatory protein *spxA* in *S. aureus* and *E. faecium*, with one homolog near a hotspot in *S. aureus*, and two homologs near hotspots in *E. faecium*. The enrichment of *hok/sok* components is interesting, and may indicate that disruption of these toxin-antitoxin systems by MGE insertion is a common adaptive strategy in both of these species (Hayes 2003).

Seven other homologous genes were found to be near multiple insertion hotspots within the same species, including three homologs of PPE family protein PPE15 in *M. tuberculosis*, two IS987 transposases in *M. tuberculosis*, Serine/threonine-protein phosphatase 1 and 2 (*pphA, pphB)* in *E. coli*, three IS5 family transposases in *A. baumannii*, two cold shock proteins cspD and cspLA in *E. faecium*, two homologs of general stress protein *glsB* in *E. faecium*, and two homologous and uncharacterized genes in *A. baumannii*. The fact that several transposases themselves are near MGE insertion hotspots suggests that these regions are already disrupted by IS insertion in the reference genome, or that IS elements are themselves frequently targeted by other MGE insertions.

Eight other homologous genes were found near hotspots across species, including lipoteichoic acid synthases *ltaS* and *ltaS1* in *S. aureus* and *E. faecium*, Phosphoenolpyruvate-protein phosphotransferase *ptsI* in *S. aureus* and *E. faecium*, ATP-dependent RNA helicase *cshA* in *S. aureus* and *E. faecium*, 6-phosphogluconate dehydrogenase *gnd* in *E. coli* and *E. faecium*, tryptophan--tRNA ligase 2 *trpS2* in *E. faecium* and *A. baumannii*, putative glutamine amidotransferase *yafJ* in *N. meningitidis* and *A. baumannii*, 50S ribosomal protein L31 type B *rpmE2* in *S. aureus* and *E. faecium*, and HTH-type transcriptional regulator *acrR* in *E. coli* and *K. pneumoniae.* Additionally, a distantly related *acrR* homolog in *A. baumannii* also overlaps an MGE insertion hotspot, indicating that disruption of this drug efflux pump repressor by MGE insertions is an adaptive strategy that is conserved in three of the nine species considered here. Considering the repeated targeting of these homologous genes by MGE insertions both within and across species, they are likely good candidate genes to investigate further for functional and adaptive significance.

An analysis of Gene Ontology (GO) enrichment of all genes near a significant insertion hotspot (FDR ≤ 0.05) identifies many pathways that are enriched near insertion hotspots (Figure 4b; Supplementary Table 8). Several of these enriched pathways are clearly associated with antibiotic resistance, including “pore complex” and “porin activity” in *K. pneumoniae*, and “drug export”, “response to drug”, “efflux transmembrane transporter activity”, and “drug transmembrane transporter activity” in *E. coli*. Other enriched pathways are associated with host infection and virulence, including “siderophore transport” in *N. meningitidis*, and “cell adhesion involved in single-species biofilm formation” in *E. coli.* Additionally, broader GO terms are enriched that generally point to transcription factor activity, such as “protein-DNA complex” and “transcription factor binding” in *P. aeruginosa*, and “DNA-binding transcription factor activity” in *N. meningitidis*. These findings suggest that MGE insertion hotspots in these populations of pathogenic isolates may specifically modulate cellular functions such as antibiotic resistance, virulence, and transcription factor activity across species.

Finally, we used HOMER motif analysis software to identify target sequence motifs for individual MGEs. We find 55 elements have significant target sequence motifs (P < 1e-11; Supplementary Table 9; Supplementary File 1). Motifs for five MGEs with particularly high target sequence specificity are highlighted in Figure 4c. We find seven elements with CTAG target-site motifs, two of which are shown in Figure 4c. This motif has been described previously for other IS elements (Fournier, Paulus, and Otten 1993). The highly specific 12 base motif identified for ISPa11 corresponds with previous studies demonstrating that this element targets repetitive extragenic palindromic (REP) sequences throughout the genome (Tobes and Pareja 2006). The target sequence specificity of the MGEs described here may be of interest to the field of genome engineering. In summary, we find that regions frequently disrupted by independent MGE insertions are associated with important biological functions such as antibiotic resistance, and that by analyzing these regions we can determine the target-site specificity of these MGEs.

### MGE insertions contribute to antibiotic resistance in laboratory evolution experiments

Adaptive laboratory evolution (ALE) experiments are powerful tools to understand how drug resistance emerges. Our approach can also be used to analyze MGE insertions in an experimental context from short-read sequencing data. We wanted to determine how frequently MGE insertions contribute to antibiotic resistance in a controlled laboratory experiment. Studies have demonstrated in laboratory grown *E. coli* that the rate of IS insertion is about 1/3rd the rate of point mutation, but it is unclear how frequently these mutations would actually affect gene function as they seemed to selectively target intergenic regions (Lee et al. 2016). Using our *mustache* insertion analysis tool, we re-analyzed data from two ALE experiments that investigated the mechanisms by which *E. coli* adapts to prolonged exposure to antibiotics. These included the megaplate experiment conducted by Baym et al. (2016), and the morbidostat experiment conducted by Toprak et al. (2011).

#### MGE insertions play an important role in megaplate-based ALE studies of antibiotic-resistance

We first analyzed the WGS data collected by Baym et al. (2016), a study that introduced the megaplate as a means of visually observing a migrating bacterial front across a landscape of varying antibiotic concentrations (Baym et al. 2016). The authors performed a series of experiments evaluating the acquisition of antibiotic resistance in the *E. coli* strain K-12 BW25113. Individual bacterial lineages were picked from the plate and shotgun sequenced in a multiple intermediate-step trimethoprim (TMP) experiment where they collected and sequenced 230 isolates. Among these sequenced isolates, we identified 34 independent IS insertions (Fig. 5a,b; Supplementary Figure 4; Supplementary Table 10). Six of the 14 insertion sequences annotated in the *E. coli* reference genome were detected at least once, including IS1, IS2, IS3, IS5, IS186, and IS30. These insertion sequences directly disrupted the coding sequence of 8 different genes: *acrR, aroK, pitA, rng, rzpR, sspA, tufA, gshA, ydjN, yeaR*, and *yeiL*. Other insertion events occurred upstream of *lon, mgrB, aroK*, and *flhD*. In this trimethoprim-resistance experiment, in comparison to the adaptive single nucleotide polymorphism (SNP) and indel mutations originally reported, insertion sequences account for 17.1% (95% CI, 11.6%-22.7%) more adaptive mutations (defined as the mutations occurring in genes that are mutated independently at least twice), and 24.2% (95% CI, 16.7%-31.7%) when *folA* substitutions/indels are excluded. This rate of IS insertion is comparable to those reported in other studies of adaptation to anaerobic conditions (Finn et al. 2017), and different growth conditions (Knöppel et al. 2018).

**Figure 5:**
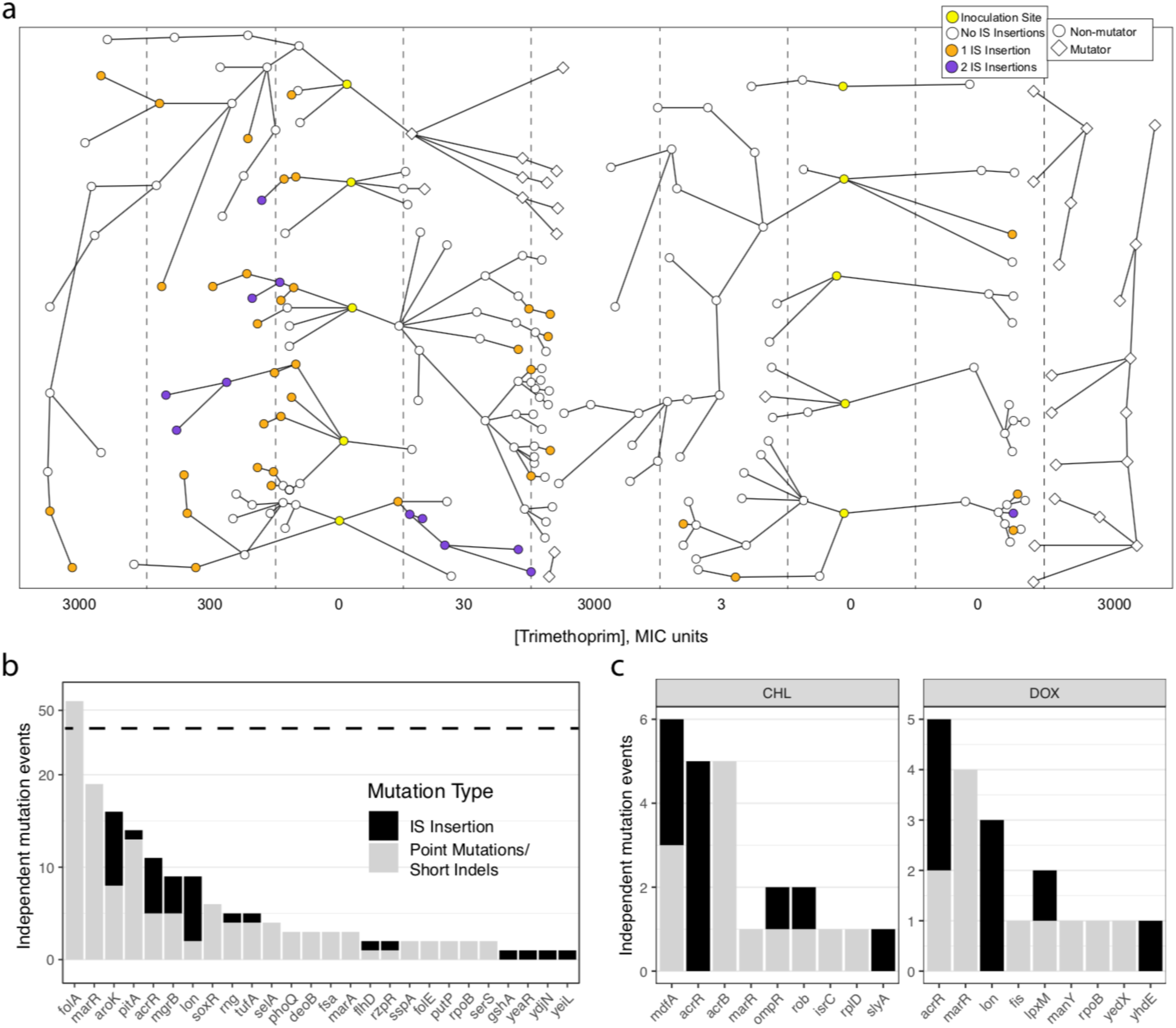
MGE insertions contribute to antibiotic resistance in adaptive laboratory evolution experiments. **a**, A schematic representation of the intermediate-step trimethoprim megaplate experiment conducted by Baym et al. (2016). Showing the number of MGE insertions found in each sequenced isolate collected from each position on the megaplate. The concentrations listed along the bottom refer to multiples of the minimum inhibitory concentration (MIC) of trimethoprim used in each panel. Apparent inconsistencies across nodes can be explained by low coverage of samples and errors when inferring descent, which was based solely on analysis of the experiment on video. **b**, The number of independent mutation events assumed to affect each gene listed in the intermediate-step megaplate experiment conducted by Baym et al. Black bars indicate the number of independent mutations caused by MGE insertion, and grey bars indicate the number of genes affected by point mutations / short indels as they were initially presented by Baym et al. This includes all newly discovered MGE insertions affecting a gene only once (*yeiL, ydjN, gshA, yeaR*) and excluding all point mutations/short indels affecting a gene only once. **c**, An analysis of the results of the morbidostat experiment conducted by Toprak et al, supplemented with IS insertion information. Showing only the results of the chloramphenicol (CHL) and doxycycline (DOX) experiments, and only showing the point mutations / short indels reported in the initial study.

Six of these target genes, *acrR, aroK, pitA, mgrB, tufA*, and *rng*, overlap with the top hits of the SNP and indel analysis performed by Baym *et al.* These were all events that either directly disrupted the coding sequence or occurred just upstream, providing strong evidence for adaptation by loss-of-function. The insertion sequences upstream of *lon* and *flhD* have been previously reported. The insertion of IS186 upstream of the *lon* gene often disrupts the promoter region (SaiSree, Reddy, and Gowrishankar 2001), leading to down-regulation of *lon*. IS insertion upstream of the *flhD* gene, a regulator of swarming motility, has also been documented (Barker, Prüss, and Matsumura 2004). This IS insertion disrupts a repressive element regulating *flhD*, leading to increased expression of the *flhD* transcription factor and its target genes. IS1 and IS5 insertions upstream of *flhD* strongly correlate with increased motility (Barker, Prüss, and Matsumura 2004). Swarming motility in general is strongly associated with antibiotic resistance (Lai, Tremblay, and Déziel 2009), suggesting that this gain-of-function insertion may be yet another trimethoprim resistance adaptation. Taken together, this gives us an estimate of the rate of adaptive mutations mediated by IS insertion in the context of TMP adaptation, explaining 17.1% more adaptive mutations than were originally reported.

#### MGE insertions play an important role in morbidostat-based ALE studies of antibiotic-resistance

Next, we analyzed the WGS data generated by Toprak et al. (2012) where they introduced the morbidostat, a selection device that continuously monitors bacterial growth and dynamically regulates drug concentrations to constantly challenge the bacterial population (Toprak et al. 2011). Toprak et al. treated drug-sensitive MG1655 *E. coli* with chloramphenicol (CHL), doxycycline (DOX), and trimethoprim (TMP), and performed whole-genome sequencing on 5 independently treated cultures from each of the three treatment groups. They identified point mutations and indels that evolved in independent lineages in response to antibiotic treatment, and reported 14, 11, and 23 such mutations for CHL, DOX, and TMP treated populations, respectively. We identified 11, 8, and 0 novel IS-mutations for CHL, DOX, and TMP treated populations, respectively (Fig. 5c). The seeming lack of IS-mediated mutations in the trimethoprim arm of this study relative to the megaplate study may indicate how the two experimental conditions may influence IS activity in different ways. When considering only the mutations originally reported by Toprak et al., in the populations treated with CHL and DOX, IS insertions caused 11/25 (44%) and 8/19 (42%) of antibiotic-resistance mutations, respectively. In the originally reported results for this study, two independent nonsense mutations were identified in the *acrR* gene across the ten CHL and DOX samples. When accounting for IS insertions, all ten CHL and DOX treated populations had a disrupted *acrR* gene, suggesting that *acrR* disruption is a major component of CHL and DOX resistance in the context of this morbidostat experiment. Additionally, the *lon* promoter-disrupting IS insertion observed in the study by Baym et al. was observed in this study in three of the five doxycycline replicates. The *lpxM* gene was disrupted in one doxycycline replicate, and *slyA* was disrupted in one chloramphenicol replicate. Again, this demonstrates the key role and high rate of IS insertions as mutations that confer antibiotic resistance, and that their inclusion in analysis of such experiments is critical to forming a more complete understanding of genetic mechanisms of adaptation.

### MGE insertions contribute to antibiotic resistance in clinical isolates

While these *in vitro* findings do suggest that MGE insertions contribute to many of the adaptive mutations to antibiotic resistance, we wanted to determine if these results were supported among clinical isolates. Our approach can help us to understand the role of MGE insertions in the acquisition of antibiotic resistance in a well-annotated clinical isolate collection. The isolates downloaded from the SRA database and described in previous sections of this study were a random sampling of what is publicly available, which limits our interpretation of the results. It is not immediately obvious what phenotypes the MGE insertion hotspots may be linked to, for example. In the case of *E. coli*, the downloaded samples may include repeatedly sequenced isolates of a few common laboratory strains, such as K-12. To address these concerns and to perform a more precise analysis, we downloaded two collections of clinical *E. coli* isolates with available antibiotic resistance phenotype information.

The first is a collection of 241 *E. coli* bacteremia isolates previously investigated in a bacterial genome-wide association study (GWAS), where the authors identified several polymorphisms, *k-*mers, and genes associated with antibiotic resistance, but did not directly investigate insertion sequence activity (Stoesser et al. 2013; Earle et al. 2016). This collection was obtained from patients at the Oxford University Hospitals NHS Trust, and will be referred to as the “Hospital” collection. The second collection includes 260 *E. coli* clinical isolates collected from various locations across the United States. Several of these isolates were sequenced and analyzed in connection with the FDA-CDC Antimicrobial Resistance Bank. This collection will be referred to as the Multi-drug resistant (MDR) collection.

Antibiotic resistance phenotypes for the Hospital collection isolates were available for the drugs ampicillin, cefazolin, ceftriaxone, cefuroxime, ciprofloxacin, gentamicin, and tobramycin (Supplementary Figure 3a). There is a range of drug susceptibilities across the isolates, with 42 isolates being susceptible to all 7 drugs tested, and 19 being resistant to all drugs (Supplementary Figure 3) tested. The MDR collection includes samples taken from a wide variety of sources, but the majority are isolated from urine (n = 146), and blood (n = 70). Isolates in the MDR collection include resistance information for 15 to 24 different antibiotics, with 16 antibiotics being tested on 200 or more samples (Supplementary Figure 3a; Supplementary Table 11). Included in the MDR collection are several isolates that are resistant to a carbapenem antibiotic (ertapenem or meropenem), a class of antibiotics often used as a drug of last resort. As expected, the Hospital collection contains more antibiotic susceptible organisms, whereas the MDR collection contains more multi-drug resistant organisms (Supplementary Figure 3b). Phylogenetic analysis indicates that isolates from both collections can be found in most major lineages (Supplementary Figure 2c,d).

We ran *mustache* on both of the isolate collections, and performed a MGE insertion hotspot analysis (Figure 6a). We compared the hotspots in each collection with the hotspots found among the randomly downloaded SRA isolates, and identified 9 hotspots that replicated across the MDR, Hospital, and randomly-downloaded SRA collections. We find that *acrR* replicated across all three collections (Figure 6a,b); hotspots near an L,D-transpeptidase, 6-phosphogluconate dehydrogenase *gnd*, and lipoprotein *yiiG* replicated between the hospital collection and the random SRA collection; and hotspots near the outer membrane porin *ompF (*Figure 6a,c), DNA-binding transcriptional activator/c-di-GMP phosphodiesterase *pdeL*, outer membrane porin C *ompC*, uncharacterized protein *ygcG*, and fructose 1,6-bisphosphatase *yggF* replicated between the MDR collection and the random SRA collection (Figure 6a,c).

**Figure 6:**
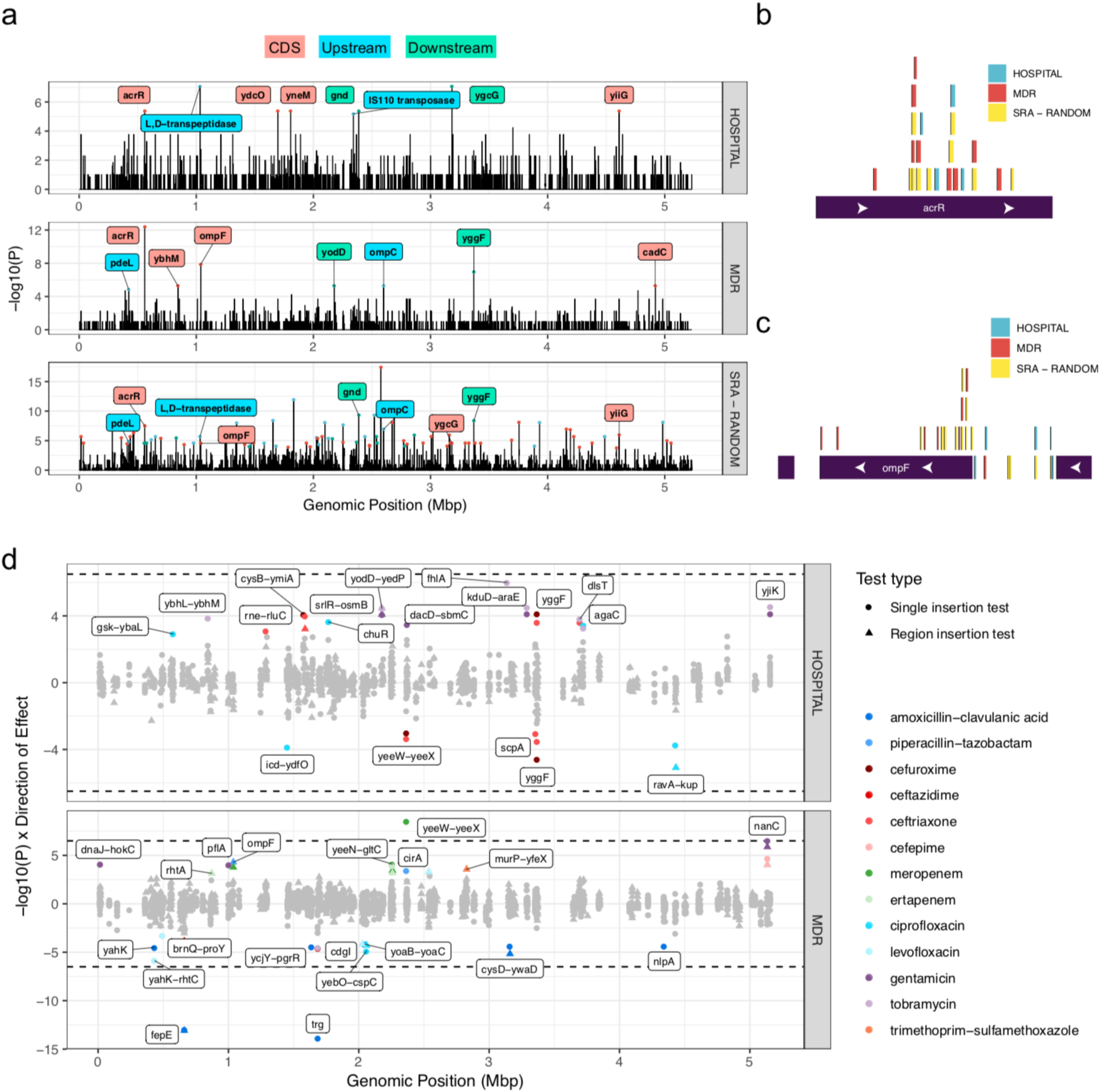
A GWAS of MGE insertions in *E. coli* identifies strong associations with antibiotic resistance in two separate isolate collections. **a**, MGE insertion hotspot analysis for the Hospital and MDR clinical *E. coli* isolate collections. The method used is the same as was used in Figure 4a for the randomly downloaded isolates. The first two panels are the hotspot analysis for the Hospital collection and the MDR collection respectively, with top hits labeled as in Figure 4a. The third panel is the same as the *E. coli* panel shown in Figure 4a but highlighting the hotspots that are shared with either of the clinical isolate collections. **b**, Showing all unique *acrR* MGE insertions found in the Hospital collection, the MDR collection, and the *E. coli* isolates downloaded at random from the SRA database. **c**, Showing all unique *ompF* MGE insertions found in the Hospital collection, the MDR collection, and the *E. coli* isolates downloaded at random from the SRA database. **d**, Results of microbial GWAS of antibiotic resistance using MGE insertions as predictors. All associations that meet a within-collection FDR-adjust P value cutoff of 0.05 are colored according to the figure legend. The y-axis is equal to the −log10(P) value of the test, multiplied by the direction of the effect, with MGE insertions that confer resistance being positive and MGE insertions conferring sensitivity being negative. Text descriptions indicate the region in which the MGE insertion has taken place: The dotted line is equal to a −log10 P value cutoff of 6.5, a stringent Bonferroni cutoff used in the comprehensive SNP, indel, and gene presence/absence GWAS originally conducted on the Hospital dataset by Earle et al. (2016).

The location of genes involved mechanisms of antimicrobial resistance near hotspots in different collections of isolates is compelling evidence that mobile element insertion hotspots indicate important adaptations, and are not merely sinks for random, functionally insignificant insertions. The *acrR* and *ompF* mutations are well-described antibiotic resistance mutations (Harder, Nikaido, and Matsuhashi 1981; Jellen-Ritter and Kern 2001a), and the impact of the mobile element insertions within the other hotpots should be investigated further for functional significance. It should also be noted that while the *gnd* hotspot is closest to the *gnd* coding sequence, it is also directly upstream of the gene *ugd*, a UDP-glucose 6-dehydrogenase, which may be the functionally relevant gene.

Finally, we wanted to determine the extent to which MGE insertions are informative in a microbial genome-wide association study (GWAS). Microbial GWAS are growing in popularity as methods for controlling for population structure in clonally reproducing species improve, and as more sequenced isolates with well-annotated phenotypes are becoming available. We used MGE insertions as predictors in a linear mixed model to predict antibiotic resistance phenotypes in the Hospital and MDR collections. For each antibiotic resistance phenotype, all resistant isolates within each collection were considered cases while the sensitive isolates were used as controls. We performed tests for the presence/absence of a single insertion at a specific site, as well as tests for the presence/absence of any insertion in a given gene or intergenic region. We used the linear mixed model implemented in GEMMA, a commonly used approach to control for population structure and sample relatedness (Zhou and Stephens 2012) (Supplementary Table 12).

We see a region between the genes *yeeW* and *yeeX* where an insertion associated with antibiotic resistance exists in both collections (MDR collection, *P* = 3e-9; Hospital collection, *P* = 4.2e-4), but with opposite directions of effect, being associated with resistance to meropenem in the MDR dataset, but sensitivity to ceftriaxone in the Hospital dataset. In the *E. coli* K-12 reference, this region contains a putative uncharacterized protein *yoeF* that was not annotated in our pipeline (Keseler et al. 2017). The insertion identified at this site is an ISEc23 (IS66 family) insertion at position AE014075.1:2368401 in the CFT073 reference genome, near the end of the cryptic prophage CP4-44. The deletion of this prophage reduces cell growth and biofilm formation (Wang et al. 2010), and it contains the YeeV-YeeU toxin-antitoxin system which has been linked to bacterial persistence. The exact impact of these insertions must be further validated experimentally, but the fact that they replicate across distinct collections, albeit in different directions, is intriguing.

Both the *fepE* coding sequence disruption (*P* = 8.2e-14), and a single IS150 insertion into methyl-accepting chemotaxis protein *trg (P* = 1.3e-14) are associated with increased sensitivity to amoxicillin-clavulanic acid. The gene *fepE* regulates LPS O-antigen chain-length, ultimately influencing virulence (G. L. Murray, Attridge, and Morona 2003). We hypothesize that coding sequence disruptions of *fepE* affect virulence in such a way that it changes the cells virulence and nature of the infection, and the subsequent antibiotics that it is exposed to. It is unclear how the *trg* disruption may be biologically related to beta-lactam/lactamase inhibitor activity. One potential model that may explain the observation that loss-of-function of the *trg* gene is associated with increased antibiotic sensitivity is that the *trg* gene product, located in the inner membrane with a region that protrudes into the periplasmic space, may serve as a sink for either amoxicillin or clavulanate, and thus may prevent antibiotic activity in the periplasm. Of course, validation of these associations and predicted mechanisms of action are necessary going forward.

Other associations can be more easily interpreted. Insertions into the *ompF* gene are associated with meropenem resistance (*P* = 5.1e-5) in the MDR collection, a gene whose disruption is a known mechanisms of resistance (Harder, Nikaido, and Matsuhashi 1981). MGE insertions into the *nanC* N-acetylneuraminic acid-inducible outer membrane channel are associated with gentamicin resistance in the MDR collection (*P* = 2.8e-7), which may reduce cell permeability to this drug. To our knowledge, the association between this particular porin and antibiotic resistance in *E. coli* has not been described previously.

In the hospital dataset, insertions into the *fhlA* gene are the strongest association, corresponding with resistance to tobramycin and gentamicin (FDR-adjusted P = 9.9e-7). This gene is a transcriptional activator for the formate hydrogenlyase system, and thus would be expected to upregulate formate metabolism (Rossmann, Sawers, and Böck 1991). Formate metabolism may increase the proton motive force (Kane et al. 2016), and it has been demonstrated that the proton motive force enhances uptake of aminoglycoside antibiotics (Allison, Brynildsen, and Collins 2011), whose target is intracellular. Thus, *fhlA* null mutants, generated by MGE insertions disrupting this gene, may confer antibiotic resistance through decreased formate metabolism and proton motive force resulting in decreased uptake of aminoglycosides such as tobramycin and gentamicin.

It should be noted that most of the associations did not meet the most stringent significance cutoff that would be used in a full microbial GWAS, and that the effects associated with these insertions are relatively small compared to the effect seen in association with the presence of a beta-lactamase gene, for example. Many of the functional effects caused by MGE insertions could also be caused by point mutations and short indels, which are not being evaluated here. But by including MGE insertions in a full microbial GWAS design, this may highlight mechanisms of resistance that would otherwise be ignored, adding additional information and improving overall predictive power. In summary, a GWAS of MGE insertions is useful for understanding genomic variation that contributes to antibiotic resistance, and we have identified several candidate genes that will require experimental validation in the future.

## Discussion

Moving beyond base-pair substitutions and small insertions/deletions will allow us to better understand genomic evolution of these important pathogenic bacteria. Here, we have demonstrated an approach to genotyping large insertions from short-read sequencing datasets that is sensitive and precise. Applying this approach to an analysis of several thousand bacterial isolates of nine different pathogenic species, we identified several known and novel MGEs. We find that the MGE mutational spectrum directly highlights genes involved in antibiotic resistance. We show that MGE insertions frequently contribute to the acquisition of antibiotic resistance in *E. coli* laboratory experiments and report the first bacterial genome-wide association study of large insertions, an analysis that highlights potential mechanisms of multidrug resistance mediated in part by MGE insertions.

MGE insertions are often ignored, as few accessible tools exist that allow them to be easily and thoroughly investigated in the context of short-read sequencing. Using an approach that allows us to thoroughly characterize MGE insertions from short-read sequencing datasets, we demonstrate that much can be learned about MGE insertions. We focus on analyzing publicly available datasets in part to demonstrate the value in re-analyzing such data with novel methods.

Our analysis provides strong evidence that as more isolates of a given species are analyzed, more MGE insertions are identified, suggesting that these elements are active in each of the species analyzed, although the level of activity of these elements seemed to vary between organisms. Of particular note, we find that certain species, such as *E. faecium*, have particularly high MGE activity; to our knowledge, this is the first time this has been shown, as estimates of the relative activity of MGEs across species are scarce and are derived from a small number of environmental isolates or laboratory experiments. This suggests that certain species rely more heavily on MGE movement as a mechanism of adaptation and evolution than other species.

Because our approach is not restricted to any one class of MGE, we identify insertion sequences, transposons, integrons, group II introns, and other classes of MGEs in this analysis. Many of the MGE insertions we identified contained passenger genes coding for known functions, such as antibiotic resistance genes; however, many MGEs contained largely uncharacterized proteins, which we anticipate may provide interesting adaptive advantages to the host bacterium. The passenger genes encoded in these MGEs can spread rapidly between organisms and selective forces likely impact the retention versus loss of these elements in individual organisms and in communities of organisms, such as microbiomes. Indeed, recent work has demonstrated that individuals with very similar gut microbiomes can harbor very different MGE repertoires (Brito et al. 2016). This suggests that monitoring the MGE potential of individual bacterial species and microbiomes will inform our understanding the extent of and consequences of MGE-derived genetic variation.

MGEs may contribute to organismal fitness by bringing new genes and functions to an organism. Alternatively, minimal MGEs such as IS elements may impact organismal fitness by insertional mutagenesis. Because our method allows us to both identify MGEs and their insertion sites, we are able to use MGE insertion sites accumulated across sequenced isolates to identify insertion hotspots throughout each bacterial genome. We find genes that are repeatedly “hit” by insertional mutagenesis, such as *acrR*, a gene involved in sensitivity to many different antibiotics (Jellen-Ritter and Kern 2001b). Insertional loss of function of the gene is identified at a high rate in *E. coli, K. pneumoniae*, and *A. baumannii* and likely correlates with increased antibiotic resistance of these organisms. In addition to identifying genes known to be involved in antibiotic resistance, we identify additional genic “hotspots” that may represent additional, novel genes involved in antibiotic sensitivity and resistance.

When we apply this method to previously published adaptive laboratory evolution experiments on *E. coli*, we find that insertions comprise a large proportion of the acquired mutations in laboratory *E. coli*. We find that by including MGE insertions in these analyses, the importance of *acrR* loss-of-function mutation as an adaptation to antibiotics is enhanced, re-prioritizing these mechanisms of resistance. A final observation that results from identifying “where” IS elements land within the genome is that certain target sites appear to be hit by specific IS elements. These sequence-specific transposases may have value in genetic engineering applications. Thus, by identifying the locations where IS elements accumulate, we can identify genes important for bacterial fitness and the molecular specificity of transposase genes.

Given the strong evidence that MGE insertions accumulate near genes involved in mechanisms of antibiotic resistance, and that MGE-mediated mutagenesis can confer antibiotic resistance in adaptive laboratory evolution experiments, it follows that MGE insertions may genetic features that can inform clinical antibiotic resistance. A GWAS analysis using IS element location as a feature demonstrates the utility of using MGE insertions as predictive features, and points to new genes and intergenic regions associated with multi-drug resistance. For example, *ompF* loss-of-function insertions and insertions in an intergenic region between *yeeW* and *yeeX* are associated with carbapenem resistance. While the mechanism associated with the *yeeW-yeeX* intergenic insertions remains unknown, the association of these findings with resistance to carbapenems, which are drugs-of-last-resort, is certainly of interest as multidrug resistance continues to spread throughout the population.

While the application of this method to clinical bacterial isolates and adaptive laboratory evolution experiments has been illuminating in identifying novel types of MGEs with and without passenger genes, genes that are potentially involved in antibiotic resistance, and the sequence specificity of novel transposase enzymes, there are several limitations of this approach. First, when attempting to identify large structural variants using only short read sequencing data, we necessarily rely on methods that require inference, which inherently limits our precision. Unless a MGE can be fully assembled in context from short-read sequencing data, we cannot say with certainty what the full inserted sequence is at a given position. MGEs that consistently fail to assemble using short-read sequencing, either in context or on their own as an individual contig, would evade detection using our approach. With the advent of more accessible read cloud and long-read sequencing approaches (Bishara et al. 2018), we anticipate that such inferences can be readily orthogonally validated. Another limitation of our approach is that it is also quite computationally-intensive, with the rate-limiting step usually being the assembly of the bacterial genome. Thus, adapting this approach for application to organisms with larger genomes may be limited by the computational requirements associated with de novo assembly. Alternatively, our approach can be used with a database of MGE queries that were previously identified, thus not requiring sequence assembly. The more comprehensive database of insertion elements generated as a part of this study can thus be leveraged in situations where accessing high performance computing resources is challenging or cost-prohibitive. Finally, although the associations that we present between gene disruption and antibiotic resistance are statistically significant, the predictions that these genes are associated with true antibiotic resistance must be orthogonally tested in proper molecular experiments that test the phenotypic consequences of gene knockout and rescue.

Leveraging knowledge linking genomic variations to clinically important phenotypes, such as antibiotic resistance, will not only improve our understanding of the biological basis of bacterial adaptation, but may also open the door to new therapeutic strategies. Indeed, recent studies with pathogens such as *Plasmodium falciparum* have found novel drug targets by leveraging genomic variations linked to drug resistance (Cowell et al. 2018). Future studies could further investigate the associations identified here. For example, disruption of the *fepE* gene is strongly associated with increased sensitivity to amoxicillin-clavulanic acid. We hypothesize that this disruption may increase cell wall permeability, as a *fepE* homolog in *Salmonella enterica* has been shown to influence O antigen polymerization in lipopolysaccharides (G. L. Murray, Attridge, and Morona 2003, 2005). The approach we outline could also be applied to metagenomic sequencing datasets with little modification to identify MGEs and their sites of insertion in a more diverse microbial community.

In conclusion, we have developed a new approach to thoroughly characterize MGEs and their sites of insertions from short-read sequencing data. We have demonstrated the utility of this approach when analyzing large publicly-available datasets, experimental data, and clinical isolates. Our analysis highlights the importance of thoroughly investigating MGE insertions in prokaryotic genomes, and that only by including these and other analysis in our workflows will we be able to understand these rapidly evolving organisms.

## Methods

### Code availability

The *mustache* command-line tool is available for download at https://github.com/durrantmm/mustache. A detailed README and test dataset are included.

### Data sources, preprocessing, and quality control

All of the data analyzed in this study were downloaded directly from public databases. The datasets are divided into three categories for clarity: 1) Randomly selected Illumina short-read datasets of bacterial isolates, 2) Short-read *E. coli* sequencing datasets generated by Baym et al. and Toprak et al. for adaptive laboratory evolution experiments (Toprak et al. 2011; Baym et al. 2016), and 3) Clinical *E. coli* isolate collections with antibiogram data. The sample selection, preprocessing, and quality control for each dataset is described below.

#### Randomly selected illumina short-read data - sample selection, preprocessing, quality control

The Sequence Read Archive (SRA) SQL database was downloaded on Sep. 25th, 2018. Potential sequencing datasets were filtered initially by available metadata to only include those samples with an estimated coverage between 50-150 per isolate, and only including samples annotated as paired-end, whole-genome sequencing samples. Reference genomes were selected by using the one provided by NCBI for each of the species of interest, which were curated by the community, with the exception of *Escherichia coli* CFT073 genome used here, which has been used as a reference for pathogenic *E. coli* strains (Palaniyandi et al. 2012; Earle et al. 2016). The reference genomes used are: *Escherichia coli* CFT073 (AE014075.1), *Pseudomonas aeruginosa* PAO1 (NC_002516.2), *Neisseria gonorrhoeae* FA 1090 (NC_002946.2), *Neisseria meningitidis* MC58 (NC_003112.2), *Staphylococcus aureus* subsp. aureus NCTC 8325 (NC_007795.1), *Enterococcus faecium* DO (NC_017960.1), *Mycobacterium tuberculosis* H37Rv (NC_000962.3), *Klebsiella pneumoniae* subsp. pneumoniae HS11286 (NC_016845.1), and *Acinetobacter baumannii* strain AB030 (NZ_CP009257.1). Additionally, we carried out all of these steps for *E. coli* using two additional reference genomes - O104:H4 strain 2011C-3493 (NC_018658.1) and K-12 substrain MG1655 (NC_000913.3) (Supplementary Figure 5).

Up to 2000 samples for each species of interest were randomly selected and downloaded. The reads in the downloaded FASTQ files were deduplicated using SuperDeduper v0.3.2 (Petersen et al. 2015) and adapter sequences were trimmed using Trim Galore v0.5.0 (Krueger 2015). Samples were then aligned to their respective reference genomes using BWA MEM v0.7.17-r1188 (H. Li and Durbin 2009). Samples were excluded if they met the following filters: median coverage less than 40, estimated average read length less than 95 or greater than 305, estimated average fragment length less than 150 and greater than 750, a FASTQC v0.11.7 “Per base sequence quality” quality control failure, a FASTQC “Per sequence GC content” quality control failure, and a FASTQC “Per base N content” failure (Andrews and Others 2010). Those sequence isolates that passed these filtering steps were analyzed further to identify MGE insertions.

#### Baym et al. and Toprak et al. datasets - sample selection, preprocessing, quality control

Baym et al. and Toprak et al. datasets were downloaded from the NCBI Sequencing Read Archive (Bioproject accessions PRJNA259288 and PRJNA274794, respectively) (Baym et al. 2016; Toprak et al. 2011). Samples were processed by removing duplicate sequences using SuperDeduper v0.3.2 (Petersen et al. 2015) and adapter sequences were trimmed using Trim Galore v0.5.0 (Krueger 2015). The Baym et al. and the Toprak et al. short-read sequences were aligned to the *E. coli* K12 U00096.2 and NC_000913.2 reference genomes, respectively, as was done in the original studies.

#### Clinical E. coli isolate collections with antibiogram data - sample selection, preprocessing, quality control

Two collections of pathogenic clinical *E. coli* isolates were used in this study. The first is referred to as the Hospital collection as it represents *E. coli* isolates collected at a single hospital, the Oxford University Hospitals NHS Trust. This included 241 isolates in total, all of which were bacteremia samples. They were downloaded from the NCBI database by querying all *E. coli* samples in BioProject PRJNA306133.

The second collection, referred in this study as the Multi-Drug Resistant (MDR) collection, includes 260 isolates collected from multiple BioProjects, including PRJNA278886, PRJNA288601, PRJNA292901, PRJNA292902, PRJNA292904, PRJNA296771, and PRJNA316321 (See Supplementary Table 1). Most of these isolates come from the projects PRJNA278886 (173 isolates, Antimicrobial Surveillance from Brigham & Women’s Hospital, Boston MA) and PRJNA288601 (52 isolates, CDC’s Emerging Infections Program (EIP)). Antibiograms were collected for samples from NCBI using the search key term “antibiogram[filter]”. This collection represents samples taken from several locations throughout the country as part of many pathogen surveillance programs. They were isolated from a variety of sources, including 70 isolated from blood, 142 isolated from urine, and 27 isolated from other sources.

Samples were processed first by removing duplicate sequences using SuperDeduper v0.3.2 (Petersen et al. 2015), and adapter sequences were trimmed using Trim Galore v0.5.0 (Krueger 2015). All isolates from both collections were then aligned to the *E. Coli* CFT073 genome (AE014075.1), a uropathogenic strain used previously as a reference for clinical isolates (Earle et al. 2016). BWA MEM was used for alignments (Kathiresan, Temanni, and Al-Ali 2014), with default settings in paired-end mode.

### MGE identification pipeline

A combination of several previously published tools and custom tools, detailed below, were used to identify MGE insertions and their sequence from short-read sequencing data. This pipeline is summarized in five steps: 1) Identifying the candidate insertion sites, 2) inferring complete sequence of inserted element, 3) inferring sequences from newly created reference insertions, 4) clustering elements and classifying them as mobile and immobile, and 5) resolving ambiguous position-cluster assignments.

#### Identifying candidate insertion sites

This step is fully implemented in the *mustache* software package under the *findflanks* command. The approach taken to identify candidate insertion sites was developed independently but is similar to one taken recently by another group who published their tool under the name *panISa (Treepong et al. 2018).* First, the alignment is parsed to identify sites where reads are clipped according to the BWA MEM alignment software, filtering out reads with a mapping quality less than 20 (-- `min_alignment_quality` parameter) or those that are clipped on both sides and have an alignment length less than 21 base pairs (`--min_alignment_inner_length` parameter). First, clipped sites where the longest clipped end falls below a total length of 8 are excluded (`--min_softclip_count` parameter). Second, clipped sites that have fewer than 4 supporting reads in total are excluded (`--min_softclip_count` parameter). Third, sites that are not within 22 base pairs of an oppositely-oriented read clipped site are excluded (`--min_distance_to_mate` parameter).

In the next step, information about the un-clipped reads overlapping each of the candidate insertion sites is calculated. This is an important quality control step that filters out small indels, which commonly cause false positives. First, sites where the ratio of the number of clipped reads to the number of total reads falls below a value of 0.15 are excluded (`--min_softclip_ratio` parameter). Next, sites where the ratio of the number of reads that have a deletion or short insertion to the total number of reads at the site exceeds 0.03 are then excluded (`--max_indel_ratio` parameter). This is repeated for deletions located at the base pairs directly adjacent to the insertion site. These filters were identified largely by iterative attempts to optimize sensitivity and precision and could be further improved in the future. The result of this step is a filtered list of candidate sites.

With this filtered list of candidate sites, consensus sequences for the flanks of the inserted element at each site are determined. It is not uncommon to see two distinct flank sequences at a single site, and an approach that could distinguish between multiple sequences at a single site was taken. The clipped ends are first added to a trie data structure. All of the unique paths from the parent node to the leaves are traversed, resulting in a list of unique sequences seen at an individual site. These sequences are then clustered with each other in a pairwise manner by truncating the longer sequence to the length of the shorter sequence, and by calculating a similarity metric as the edit distance divided by the total length of the shorter sequence. This matrix of similarity scores is then analyzed to identify all connected components, with connections existing between all sequences with a similarity greater than 0.75. Each component of sequences forms its own cluster, and this cluster is then analyzed to determine a consensus sequence.

Next, a consensus sequence for a cluster of sequences is constructed by traversing down the trie data structure and taking the base pair with the highest average quality score at each level of the trie as the consensus. The trie structure is traversed until only one base read supports a given site, and the consensus sequence terminated at this point. Consensus sequences that fall below 8 base pairs in length (`--min_softclip_count` parameter) and that have fewer than 4 clipped ends supporting their existence (`--min_softclip_count` parameter) are excluded. Finally, the list of candidate sites is filtered again to only include sites that are not within 22 base pairs of an oppositely-oriented read clipped site (`--min_distance_to_mate` parameter).

These candidate sites are then paired with other sites that within a specified distance and oriented in the opposite direction using the *mustache* command *pairflanks*. These oppositely-oriented flanks are allowed to be up to 20 bases away from each other, as many MGEs create a direct-repeat upon insertion. Since many MGEs, particularly IS elements, have inverted repeats at their ends, flanks that share inverted repeats near their termini are prioritized. Ties are then broken first by pairing flanks that have similar number of supporting clipped reads, and then by the difference the length of the consensus sequences. If ties still exist, the pairs are randomly assigned to each other, but this is a rare occurrence. If no inverted termini exist between any of the pairs, then only the number of clipped reads and the difference in consensus sequence length are used to pair flanks.

For the randomly downloaded SRA samples, a minimum supporting read count of 4 for each identified flank was required. The Baym et al. data included several isolates that were sequenced at low coverage (less than 20). For these low-coverage samples, a minimum supporting paired read count of 2 (`--min_softclip_count` parameter) was required, which should be roughly as sensitive as the singled ended read count filter of 4 used in their initial study (2 at each clipped site, for a total of 4 in the pair), the consensus sequence at each site was built from all available clipped reads by setting the `-- min_count_consensus` parameter to 1, and a minimum consensus length of 4 was used (`-- min_softclip_length` parameter). For Toprak et al., and clinical *E. coli* isolates with antibiogram data, a minimum supporting read count of 4 (`--min_softclip_count` parameter) was used.

#### Inferring the complete sequence of inserted elements

Once candidate read flank pairs have been identified, the next step is to infer the full inserted element. This approach combines a variety of methods of inferring the identity of each insertion. The approach taken is described here in detail.

First, the identity of the inserted sequence is inferred from the assembly of the sequenced isolate using the *inferseq-assembly* command in *mustache*. For each pair of flanks identified, each flank is aligned to the assembly using 25 bases of context sequence for each flank in single-end mode (Supplementary Figure 1a). If these consensus flanks with 25 base pairs of context sequence align to the assembled isolate with the proper orientation, one can assume with high confidence that the intervening sequence is the complete inserted sequence. This sequence is described as “inferred from assembly with full context”. If only one of the flanks align with context, and the other partially aligns to the edge of the assembled contig, this sequence is described as “inferred from assembly with half context”. These are the highest quality inferred sequences, and they are prioritized above all others when genotyping an insertion.

Next, flanks are aligned to the assembly without any context sequence. Often, these flanks align to small assembled fragments, with short parts of the flank ends being clipped off at the ends of the assembled contig. The full sequence is inferred by including the clipped flank ends, and the full intervening sequence (Supplementary Figure 1b).

The next approach to infer the sequence identity is implemented in the *inferseq-overlap* command of *mustache*. This infers the identity of the sequence by attempting to find high-confidence overlaps between the two flank ends. This can only identify inserted elements that can be spanned by the two flanks, which limits its usefulness in this study where most insertions of interest are larger than 300 base pairs. It is, however, useful for filtering out smaller insertions that are found to be below this 300 base pair length limit.

Next, flanks are aligned to the reference genome using the *inferseq-reference* command, and inserted sequences are inferred in a similar manner to those inferred from assemblies without sequence context (Supplementary Figure 1c).

At each of these sequence inference steps, all candidate insertions where the alignment scores of both flanks are equally high are returned. For example, if a given IS element is found in multiple locations throughout the reference genome, all of these locations will be reported as inferred elements (Supplementary Figure 1e).

#### Inferring sequences from dynamically created reference insertion database

For the purposes of this study, a filter was applied to all of the candidate insertions to only include those predicted to be between 300 base pairs (bp) and 10 kbp in length. This database of candidate insertions is filtered to reduce redundancy, and these inferred elements were then clustered across all isolates for a given species at 99 percent nucleotide similarity using CD-HIT-EST (W. Li and Godzik 2006; Fu et al. 2012). Any sequence that was inferred from the assembly, by flank overlap or from the reference, is included when constructing this finalized database. This gives us a database of elements that are then used as a final reference database to infer insertion sequences across all samples for a given species.

This final sequence inference approach, implemented in the *mustache* package as the *inferseq-database* command, is similar to the sequence inference approaches implemented in the *mustache* commands *inferseq-assembly* and *inferseq-reference*, but with a default minimum percent identity of the aligned flanks of 90% (as opposed to the the 95% minimum identity required for other inference commands), and the requirement that the aligned flanks map within 10 bases of the ends of each element in the database in order to be considered a candidate for the inserted element at a given position (`-- max_edge_distanc`e parameter). This serves as the most sensitive inference approach, but the results depend on how the query database is constructed.

#### Clustering elements and assigning initial genotypes

At this point in the pipeline, thousands of insertion positions have been identified, and the exact identity of those insertions could be any number of hundreds of different inferred sequence elements. Much of this information is surely redundant, however, as the differences between elements may amount to one or two single nucleotide variations. To collapse this small level of heterogeneity, sequences were clustered using CD-HIT at 90% similarity across 85% of their sequence (W. Li and Godzik 2006; Fu et al. 2012). For example, if elements X and Y are found at position A in the reference genome in two different isolates, it is assumed that elements X and Y are in essence the same sequence if they cluster at 90% similarity across 85% of their sequence, and it is assumed they are completely different elements otherwise.

Before proceeding, an additional quality control step was taken. The *mustache* algorithm itself was initially run with a maximum inferred element size cutoff of 500 kilobase pairs, and a minimum size cutoff of 1 base pair. In some cases, the identity of a given insertion included several elements of differing lengths, some of those elements being outside the size range of our initial 300 base pair to 10 kilobase pair filter. This information was used to filter out elements that had high-confidence inferred sequence identities that existed outside of the 300 base pair to 10 kilobase pair range. The tools in this study are not optimized to identify insertions outside these size ranges and excluding them makes the results more reliable.

Once elements were filtered by size and organized into clusters, insertions are assigned to a final genotype as follows: First, if a given insertion is inferred from the sequence assembly with full sequence context it is given highest priority and the position is assigned to that element’s cluster. Second, if an insertion is inferred from the assembly with half context it is assigned to that element’s cluster. Third, if an insertion is inferred by flank overlap, it is assigned to that element’s cluster. Fourth, if an insertion is inferred from the assembly without context, the position is assigned to that element’s cluster. Finally, if an insertion is inferred from the reference genome or from the aggregated insertion database, the position is assigned to their respective clusters. It should be noted that the strand orientation of the inserted element was ignored in this analysis. For example, if element X was inserted at position A in isolate 1, and element X was also inserted at position A in isolate 2 but with the reverse orientation, this will be considered an identical genotype for the purposes of this study.

#### Resolving ambiguous position-cluster assignments

In the next step, we seek to resolve ambiguous position-cluster assignments. Even after clustering elements and assigning clusters to insertion sites in this manner, many ambiguities may exist. If the ends of two different elements are very similar to each other, and yet they do not cluster together at the 90% similarity threshold, a given insertion may have mapped to both of these clusters. It is important that these ambiguities are sufficiently resolved for many of the analyses conducted in this study. We took the following steps to increase our confidence in the identity of these inferred elements, and these steps were implemented primarily in custom R scripts.

First, the number of non-ambiguous positions within the reference genome where each element cluster is found are counted. This is first done for only insertions that are inferred from the isolate’s assembly with sequence context, the highest confidence set. If a given cluster is found at more than one of these high-confidence insertion positions, it is classified as a “strict MGE”. These requirements are then relaxed to include all non-ambiguous position-cluster assignments, the next highest confidence set, and all clusters that are found at more than one position in the reference genome are classified as “lenient MGEs”.

Next, ambiguous element assignments are counted. The ambiguous element assignments are first resolved by calculating the frequency of each cluster at each position within our population and assigning the ambiguous insertion to the most frequent cluster at that position. For example, if an ambiguous insertion at position A in isolate 1 maps to both clusters X and Y, and throughout the population cluster X is found more frequently than cluster Y at position A, then cluster X is assumed to be the correct assignment at position A in isolate 1.

Finally, if any ambiguous position-cluster assignments remain, they are then resolved by prioritizing “strict MGEs” described above. Our assumption is that if the insertion at position A maps to both clusters X and Y, and X is a MGE whereas Y is not, then assign position A is assigned to cluster X. If any ambiguous position-cluster assignments remain after this step, clusters are assigned using the “lenient MGEs”. If a given insertion is still ambiguous, it is left in as an ambiguous insertion, but still considered in several analyses where cluster ambiguity is irrelevant.

#### Final quality control filter to remove low-confidence elements

As a final quality control measure, elements that were never successfully inferred from the sequence assembly of any analyzed isolate were removed. When analyzing the results from the randomly selected Illumina short-read data, it was found that 306 element clusters (9.3%) corresponding to 130 unique insertion sites (0.34%) were only identified by inferring their identity from the reference genome (See Supplementary Figure 1c). That is, these elements were never identified within their expected sequence context within the assembly, and they were never identified outside of the expected sequence context within the assembly. We considered these to be low-confidence elements, and both the elements and their sites of insertion were excluded from all subsequent analyses.

### Simulations of key steps in the pipeline

To test the sensitivity and precision of our pipeline, we carried out various simulations. Here we demonstrate that the candidate insertion identification and sequence inference steps are both sensitive and precise. The organisms simulated and the accession numbers of their genomes are as follows: *Escherichia coli* CFT073 (AE014075.1), *Mycobacterium tuberculosis* H37Rv (NC_000962), *Pseudomonas aeruginosa* PAO1 (NC_002516), *Staphylococcus aureus* NCTC8325 (NC_007795), *Vibrio cholerae* O1 biovar El Tor strain N16961 (NC_002505 and NC_002506b) (Treepong et al. 2018). For each genome, the mutation rate, coverage, and library read length were combinatorially selected and ten replicates of each combination of parameters were carried out. We simulated 32 MGE insertions by randomly selecting insertion sites, using 16 species-specific IS elements downloaded from ISfinder and chosen at random, and 16 randomly generated sequences between 300 and 10,000 base pairs in length. We simulated two mutation rates (0.001 mutations per base pair and 0.0085 mutations per base pair, with 10% of mutations being indels) using DWGSIM v.0.1.11-3 (https://github.com/nh13/DWGSIM). One study of *S. aureus* strains found the average pairwise distance between strains to be 0.0085 (Takuno et al. 2012), which motivated this choice of mutation rate. Theses simulations included various genome coverages (5, 10, 20, 40, 60, 80, and 100x), and various read lengths (100, 150, 300bp), and aligned reads to genomes in paired-end mode with BWA MEM (Kathiresan, Temanni, and Al-Ali 2014). In total, this resulted in 420 simulated genomes per species. ANOVA tests implemented in R were used to analyze which factors influence precision and sensitivity. The *mustache* tool was run on these simulated samples with the default parameters.

For comparison, these simulated genomes were also analyzed with *panISa*, and the results were compared directly to those of *mustache*. The parameters used for *panISa* were chosen to most closely reflect the *mustache* default parameters and included setting the minimum number of clipped reads to consider an IS insertion to 4 using the `--minimun` parameter.

In Figure 1a, the sensitivity and precision curves shown are based on whether or not the recovered flanks are found near the expected insertion site, with flanks being considered true positives if they are 90% similar to the flanks of the true inserted elements. Similarity here is calculated as the edit distance divided by the length of the flank. Additionally, several of the other sequence inference techniques available in *mustache* were tested to determine their sensitivity (Supplementary Figure 6).

### Unique insertions accumulation curve

A unique insertion is defined as a specific cluster assigned to a specific insertion position. Using the insertions that were identified, the number of new insertions identified as additional samples were analyzed was calculated. The ‘specaccum’ function provided by the vegan package v2.5-3 was used to estimate the accumulation curve for these insertions (Oksanen et al. 2011). The “random” method was used to estimate the curve, with 100 permutations.

### Calculating rare insertions per megabase

For each species, the number of rare insertions per megabase of genome was calculated. The allele frequency of each insertion was calculated by dividing the number of isolates where the insertion is observed by the total number of isolates analyzed for the species. This list was then filtered to only include insertions with an allele frequency less than 0.01 (1% of isolates), and we then calculated the number of such rare insertions per isolate. The number of megabases for each sample with non-zero read coverage was then calculated by analyzing the alignment files for each isolate, and the number of rare variants per isolate was divided by this number of megabases.

### Annotating identified elements

All identified elements were annotated using the Prokka v1.13 annotation software (Torsten Seemann 2014). This approach uses a rapid hierarchical approach to classify proteins, with the databases being derived from UniProtKB. Default settings were used, with the exception of the added flags ‘-- kingdom Bacteria’ and ‘--metagenome’. The ‘metagenome’ flag was used to improve prediction of genes in short contigs. All unique identified elements were annotated, not just the cluster representatives.

Transposases are poorly annotated using the Prokka annotation pipeline, and they are often described only as “hypothetical” proteins, so a different approach to identify potential transposases was necessary. The profile hidden markov models (pHMMs) constructed by the creators of the *ISEScan* software were used to identify potential transposases (Xie and Tang 2017), using Prokka predicted proteins as inputs. All such proteins with *e* value < 10^−4^ were considered to be transposases. These predicted transposases were used as a reference for constructing Figure 3. Elements that contained only predicted transposases and no other ORFs were classified as IS elements (Figure 1d).

### Describing MGEs with unannotated passenger genes

An unannotated gene is defined as one that was only described as “hypothetical” by Prokka and was not annotated as a transposase using the *ISEScan* profile hidden markov model. A single MGE cluster may not always be annotated in the same way across all members of the cluster, as mutations may disrupt open-reading frames in some members of the cluster. To account for this, the number of unannotated genes in each member of the cluster was counted, and then the average number of annotated genes across all members was calculated. This number was rounded to the nearest whole number, and this value was considered to be the predicted number of unannotated genes contained within the MGE. Figure 3b shows all of the MGEs that contain two or more unannotated genes.

### Describing MGEs with annotated passenger genes

Figure 3c was generated to summarize the annotated passenger genes that were contained within predicted MGEs. If any member of a given element cluster was found to contain an annotated gene, then the entire cluster was predicted to contain the identified gene. The number of insertion sites where the predicted MGE at the specified site contained the annotated gene of interest was calculated. The passenger genes that appeared at the most unique locations throughout each organism’s genome are presented in Figure 3c.

### Annotating reference genomes

Rather than relying on the annotations provided by the NCBI GFF, each organism’s reference genome was annotated using Prokka to ensure that annotations across species were comparable to each other in quality. Prokka v1.13 was used with default settings. The gene names identified by Prokka were used to describe genes in Figures 4a and 6a (Supplementary Table 6).

### Insertion hotspot analysis

Regions of the genome with high numbers of unique insertions are described in this study as insertion hotspots. The approach taken here resembles approaches taken by ChIP-seq peak calling algorithms, such as MACS (Feng et al. 2012). All unique insertions found throughout each organism’s genomes were extracted. Sliding windows of 500 base pairs, with 50 base pair steps, were created across each organism’s genome. The number of insertions found within each window was calculated using bedtools (Quinlan and Hall 2010). To determine if a given window was enriched for insertions, a Poisson distribution was used to model the insertion distribution, with a dynamic parameter *λ*_*local*_. This parameter is calculated as

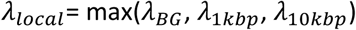

Where the term *λ*_*BG*_describes the estimated rate of insertion calculated across the entire genome, *λ*_*1kbp*_ is the estimated rate of insertion within 1 kbp of the window being tested (500 bases on either side of the window), and *λ*_*10kbp*_describes the estimated rate of insertion within 10 kbp of the window being tested. The estimated value of *λ*_*local*_ is then used in a one-sided exact Poisson test to determine if the observed insertion rate for the window exceeds the expectation, using the ‘poisson.test’ function in R. Adjusted for the local insertion rate, this should account for biases in the local insertion rate, identifying windows that are significant above this background level.

Significant insertion hotspots were then calculated as those that met FDR≤0.05. Once these significant insertion hotspots were identified, overlapping hotspots were merged into single regions. These insertion hotspots were then associated with nearby genes. This was done by first restricting the edges of each hotspot to begin and end at the exact site of the first and last insertions that they contain, effectively tightening the region edges to directly surround their insertions. The center point of each region was calculated and used to associate this region with the nearest gene. This resulted in a list of significant hotspots associated with specific genes in the reference genome.

### Insertion hotspots near homologous genes within and across species

To determine if homologous genes within and across species were found near MGE hotspots, all protein sequences that were mapped to insertion hotspots previously were extracted. These proteins were then clustered using the CD-HIT algorithm at 40% sequence identity across 70 % of the sequence. If protein sequences clustered with each other across species, they were considered cross-species insertion hotspots. If multiple protein sequences near hotspots from the same species clustered with each, they were considered within-species insertion hotspots.

### Insertion hotspot gene ontology enrichment analysis

This approach was taken to determine if the genes near hotspots were enriched for any particular function. Gene ontology (GO) terms were used for this purpose. To map genes to gene ontology terms, the Prokka predicted protein sequences were mapped to the UniRef90 database (Suzek et al. 2015) with DIAMOND (Buchfink, Xie, and Huson 2015) with an *e* value cutoff of 10^−5^, choosing the top result as the representative. The database identifier mapping (‘Retrieve/ID mapping’) service provided by UniProt was used to map the IDs of each protein to all other proteins that clustered at the level of UniRef90, and then mapped these proteins to the GO terms associated with them in the UniProt database. In other words, the GO terms associated with each annotated protein include all of those assigned to it or any homologs that cluster with it in the UniRef90 database. All of the coding sequences near an insertion hotspot were taken, and enriched GO terms were identified using the hypergeometric test, with all proteins with at least one GO term of any type used as a background. Only GO terms containing five or more genes, with two or more of these genes being found near a hotspot, were tested. A significance cutoff of FDR ≤ 0.05 was used to determine significant GO enrichments.

### Target-sequence motif discovery

The HOMER motif analysis tool was used to identify target-sequence motifs for MGEs with 10 or more different insertion sites. HOMER findMotifsGenome.pl was executed with default parameters, and searched for motifs of size 4, 6, 8, 10, and 12 within the reference of genome of each species. Randomly selected sequences from the reference genome were used as background sequences. All *de novo* motifs that met a p-value cutoff of 1e-12 were reported, choosing the most significant motif for each MGE.

### Analysis of Baym et al. megaplate experiment sequencing data

The large insertions found in the sequenced isolates from the intermediate-step trimethoprim megaplate experiment conducted by Baym et al. were analyzed. This included 231 sequenced isolates in total; of note, the sequenced isolates had widely varying sequence coverage. The sample sequencing coverage was calculated as the median sequencing coverage across the entire genome. Across all 231 isolates, the median isolate was covered by 19 reads, with 31% of isolates having median coverage less than 15, and 5.6% being covered at a genome-wide median coverage of 0. As was done in the original publication, all sequenced isolates were analyzed, including these low-coverage samples to see if any insertions could be identified, but it should be noted that due to low coverage not all insertions in all samples could be identified. This range of sequencing coverage can explain some of the unexpected patterns of inheritance observed in Figure 5a.

The samples in this experiment were analyzed with the complete insertion identification workflow. If an isolate was found to have a median coverage lower than 20, more lenient parameters were used by *mustache* to identify insertions (See ***Identifying candidate insertion sites***).

Baym et al. inferred the pattern of inheritance from watching a video recording of the migrating bacterial front as it grew across the megaplate. Their visually-inferred phylogeny was used to determine how many independent insertions occurred across all sequenced samples in this re-analysis of their data. In most cases, the inferred relationships between isolates was accurate; however, in a subset of cases, it appears that the inferred relationships between isolates was inaccurate. Thus, rather than relying completely on the given phylogeny, the number of independent insertions was estimated conservatively. An illustration of the independent insertion events that were identified is provided in Supplementary Figure 4.

### Analysis of Toprak et al. morbidostat experiment sequencing data

The trimethoprim, chloramphenicol, and doxycycline morbidostat experiments conducted by Toprak et al. were analyzed in this study. This included 20 samples in total, with 5 replicates per experiment, the wild type reference, and four additional samples that represented four additional colonies that grew from plating three of the replicates. All samples were covered at a median coverage between 21 and 30, suggesting that there was sufficient sensitivity to detect most insertions in all samples. These samples were sequenced using a single-end sequencing approach, which *mustache* is able accommodate with some minor changes in the workflow.

These samples were analyzed using the same workflow used in other analysis, with a minimum clipped read count of 4 to support each insertion site. Independent mutations were identified and cross-referenced them with genome annotations to determine the genes that were most likely impacted by each insertion. It should be noted that Toprak et al. were very stringent when initially calling SNPs and indels from their datasets, requiring validation by Sanger sequencing. The IS insertions found in this study were not subject to the same strict standards of evidence.

### Genome-wide association study of antibiotic resistance in *E. coli*

A genome-wide association study (GWAS) was implemented using only the insertions identified by *mustache* as input features. Sites that were considered to be missing (due to low coverage, defined as 10 read counts or less for either the 5’ or 3’ end of the insertion) in greater than 5% of samples were excluded. A feature matrix was then constructed with two types of predictors - the presence/absence of a specific element at a specific site and the presence/absence of any insertion within an intergenic region or coding sequence. In the Hospital collection, antibiotic resistance phenotypes were collected from a previous publication (Treepong et al. 2018). For the MDR collection, not all isolates were tested against all antibiotics. Only phenotypes where more than 200 samples were tested against the antibiotic of interest and more than 40 isolates were found to be resistant were analyzed. For both collections, isolate relatedness was calculated using GEMMA from the core-genome biallelic SNPs identified by the SNIPPY pipeline (T. Seemann 2015). A linear mixed model was used to perform association tests in GEMMA, and LRT p-values for each tested gene were kept. In the Hospital collection, no additional covariates were included in the model. In the MDR collection, isolation source was used as a covariate (blood, urine, or other). All tests that met a significance cutoff of FDR ≤ 0.05 were considered significant signals for the purposes of this study. A more stringent cutoff that is more reflective of a complete microbial GWAS (Earle et al. 2016) is shown in Figure 6d by a horizontal dotted line.

## Acknowledgments

We thank Duncan MacCannell at the Center for Disease Control (CDC) for his assistance identifying clinical isolates to analyze in this study, Christopher T. Walsh at Stanford University and Kaitlin Tagg at the CDC for their helpful feedback and critiques, Stephen Montgomery and Brunilda Balliu and other members of the Montgomery Lab for their feedback and guidance, and Eli Moss and other members of the Bhatt Lab for their feedback and guidance. This work was supported in part by the NIH grant P30 CA124435 which supports the following Stanford Cancer Institute Shared Resource: the Genetics Bioinformatics Service Center. This work was partially supported by a Donald E. and Delia B. Baxter Foundation Faculty Scholar award to ASB, and the National Science Foundation Graduate Research Fellowship to MGD.

