## Supplementary Figures and Materials for "Mobile genetic element insertions drive antibiotic resistance across pathogens"

### Supplement

**Supplementary Figure 1: Schematic representations of important concepts in the *mustache* workflow.**

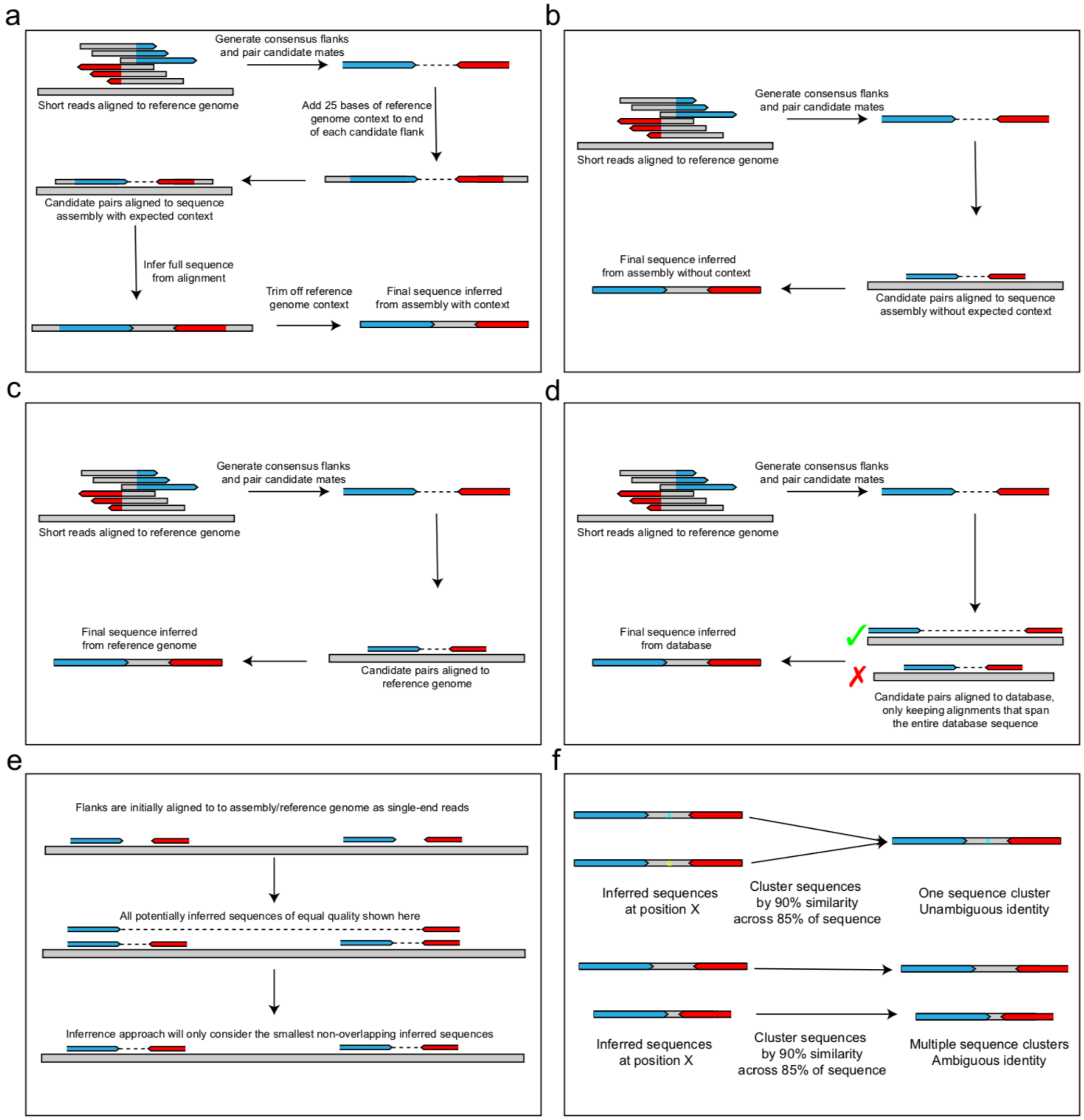


**a**, A schematic of what is meant by "Inferred from assembly with full context". "Inferred from assembly with half context" is similar, but the inferred sequence resides at the edge of an assembled contig, so only the context of one end is known with certainty. **b**, A schematic of what is meant by "Inferred from assembly without context". **c**, A schematic of what is meant by "Inferred from reference genome". In Figure 1c, these are grouped together under the label "Inferred from dynamically-constructed database". **d**, A schematic of what is meant by "Inferred from dynamically-constructed database". **e**, An explanation of how sequences are inferred from reference genomes and assemblies. In some cases, insertion sequences may occur in tandem in the reference. It is possible that the sequence that has been inserted is actually the two tandem insertion sequences and the intervening sequence, but in our workflow, we ignore this possibility and assume the smallest nonoverlapping inferred sequences to be the most accurate. **f**, A schematic of an inferred sequence with an unambiguous and an ambiguous identity. Methods for resolving inferred sequences of ambiguous identity are described in the methods.

**Supplementary Figure 2: Comparison of *mustache* and *panISa* insertion identification tools.**

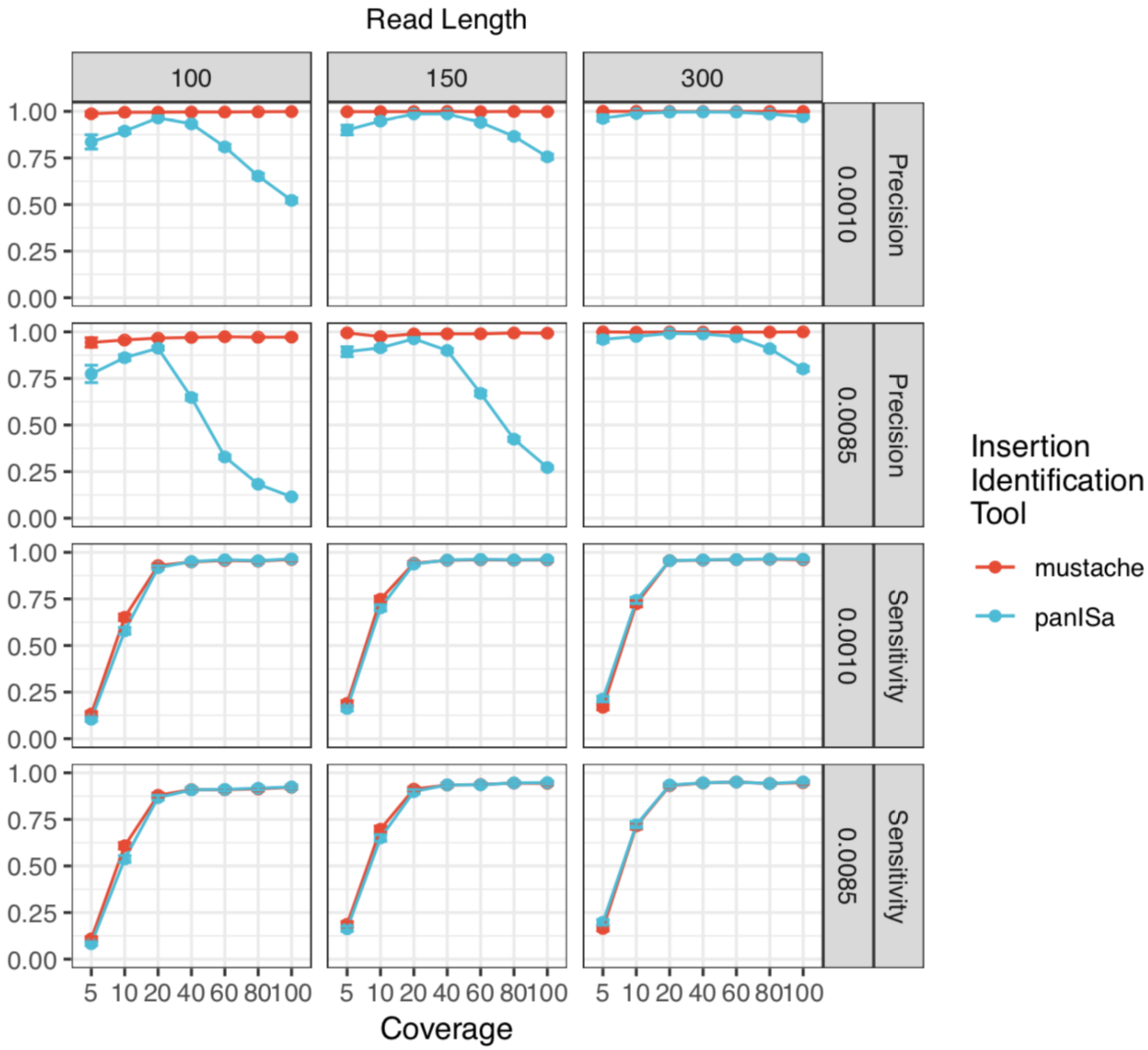


A comparison of precision and sensitivity of *mustache* and *panISa* insertion calling pipelines. Separating comparisons by read length (100, 150, 300), coverage (5, 10, 20, 40, 60, 80, 100), and mutation rate (0.0010 mutations per base pair, 0.0085 mutations per base pair). Error bars refer to 95% confidence interval of each measurement. Measurements are averaged across the simulated genomes of the 9 species investigated in this study.

**Supplementary Figure 3: Summary of two publicly available collections of pathogenic *E. coli* isolates analyzed in this study.**

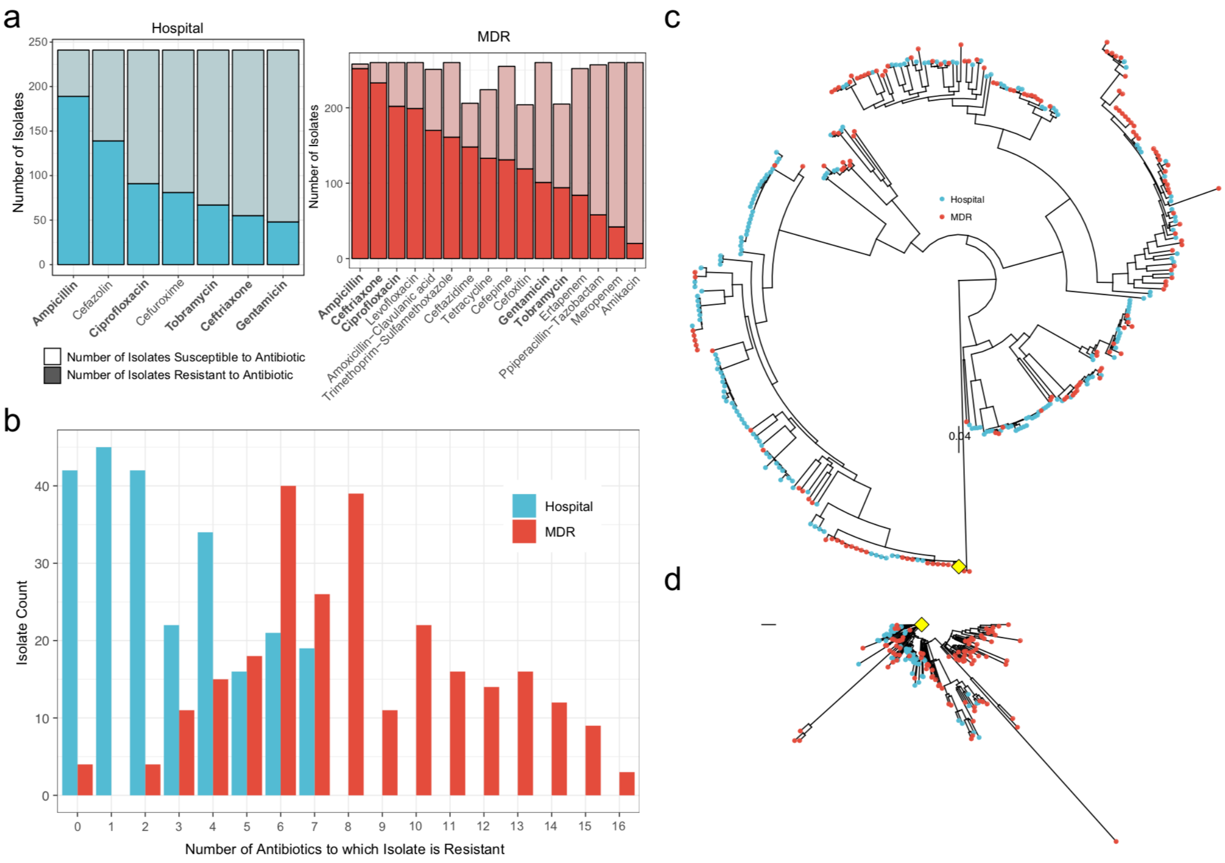


**a**, The number of samples resistant to seven different antibiotics tested in the Hospital collection (left panel, blue), and the number of samples resistant to 16 different antibiotics in the MDR collection (right panel, red). In these bar plots, darker shades indicate the total number of isolates that are resistant to the antibiotic, and lighter shades indicate the total number that are sensitive. Only those antibiotics that were tested on 200 or more isolates in the collection are shown in the right panel. **b**, The total number of antibiotics to which a sample is resistant in the Hospital collection (left panel), and the MDR collection (right panel). Only the antibiotics shown in **a,** are considered in the counts shown in both panels. **c**, A phylogenetic tree of all the samples analyzed among the two collections. Red dots indicate samples that are found in the MDR collection, and blue dots indicate the samples found in the Hospital collection. The blue diamond indicates a truncated lineage, expanded in **d**. **d**, The lineage indicated in **c** by the blue diamond, expanded here for improved visualization.

**Supplementary Figure 4: Identifying independent insertion events from the intermediate-step trimethoprim megaplate experiment.**

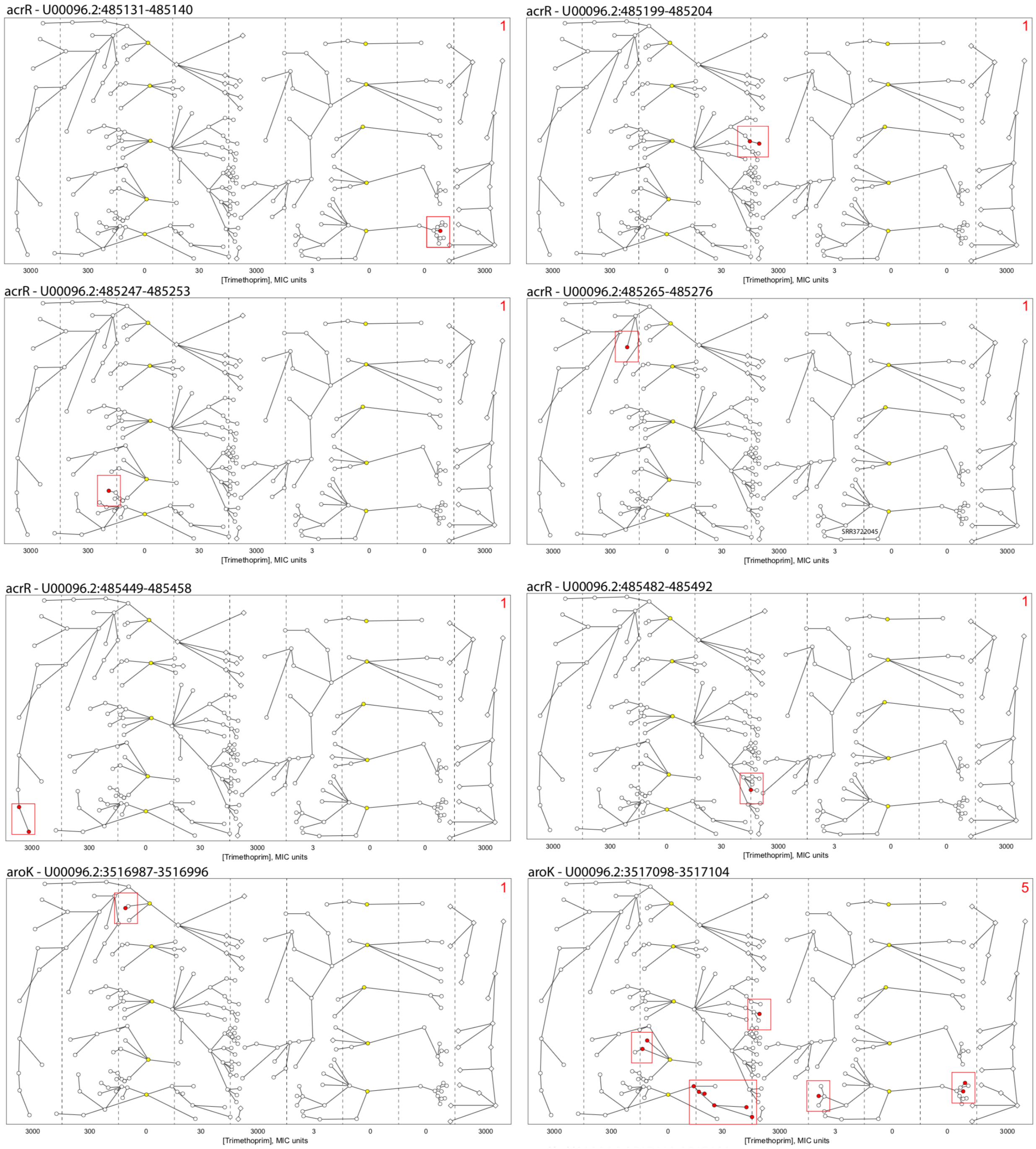

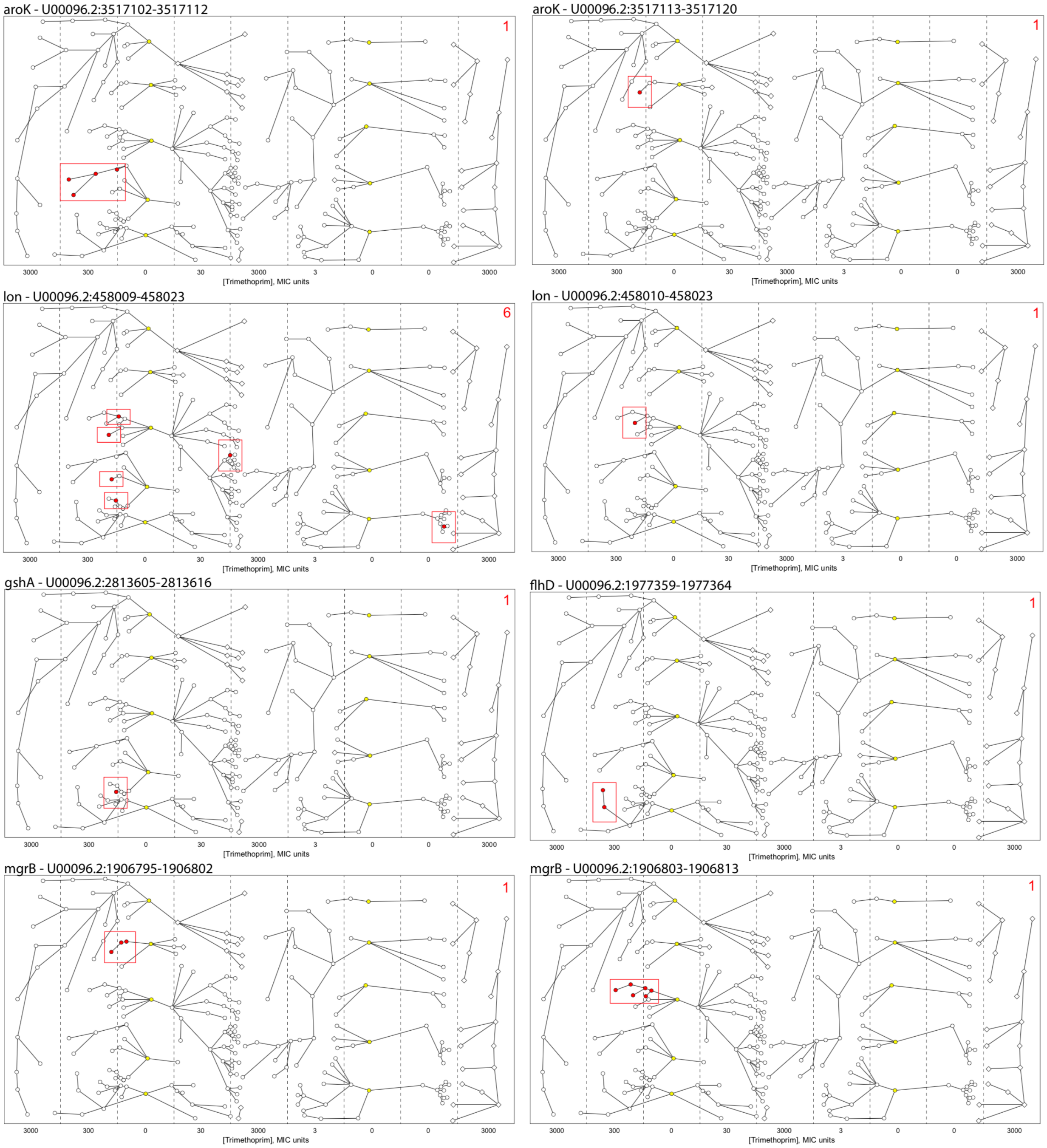
aroK - U00096.2:3517102-3517112
1
3000 300 0 30 3000 3 0 0 3000
[Trimethoprim], MIC units
aroK - U00096.2:3517113-3517120
1
3000 300 0 30 3000 3 0 0 3000
[Trimethoprim], MIC units
lon - U00096.2:458009-458023
6
3000 300 0 30 3000 3 0 0 3000
[Trimethoprim], MIC units
lon - U00096.2:458010-458023
1
3000 300 0 30 3000 3 0 0 3000
[Trimethoprim], MIC units
gshA - U00096.2:2813605-2813616
1
3000 300 0 30 3000 3 0 0 3000
[Trimethoprim], MIC units
flhD - U00096.2:1977359-1977364
1
3000 300 0 30 3000 3 0 0 3000
[Trimethoprim], MIC units
mgrB - U00096.2:1906795-1906802
1
3000 300 0 30 3000 3 0 0 3000
[Trimethoprim], MIC units
mgrB - U00096.2:1906803-1906813
1
3000 300 0 30 3000 3 0 0 3000
[Trimethoprim], MIC units

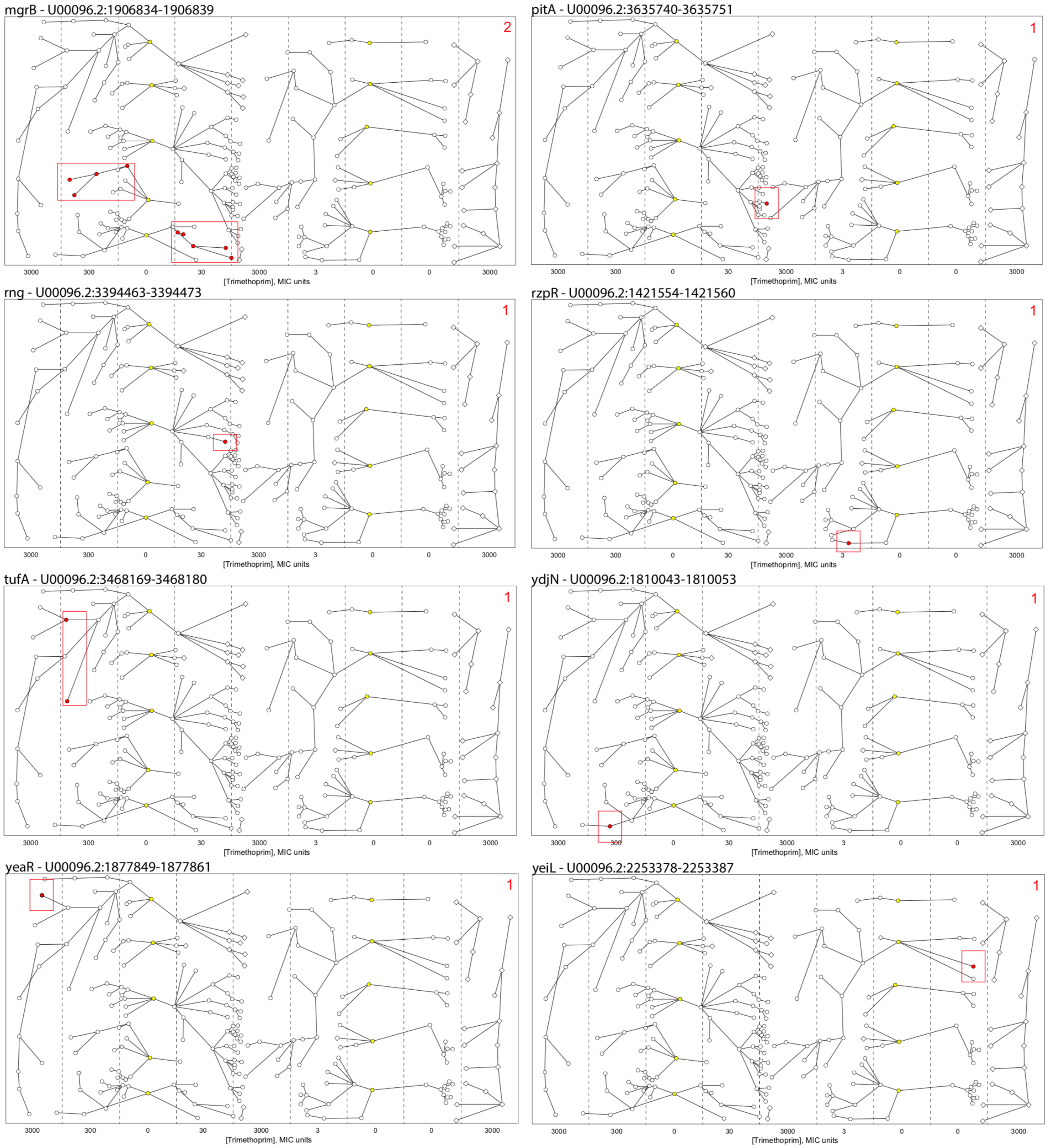


Each panel indicates how independent IS insertions were counted in the intermediate-step trimethoprim megaplate experiment conducted by Baym et al. (Baym et al. 2016) The relevant mutation is described in the top left corner of each panel, with the relevant gene named, followed by the genomic location of the insertion. Red points indicate isolates where the relevant mutation was found. The red boxes highlight a group of mutated isolates that are presumed to share a common ancestor where the initial mutation took place.

**Supplementary Figure 5: Comparison of difference between results when using three different *E. coli* reference genomes.**

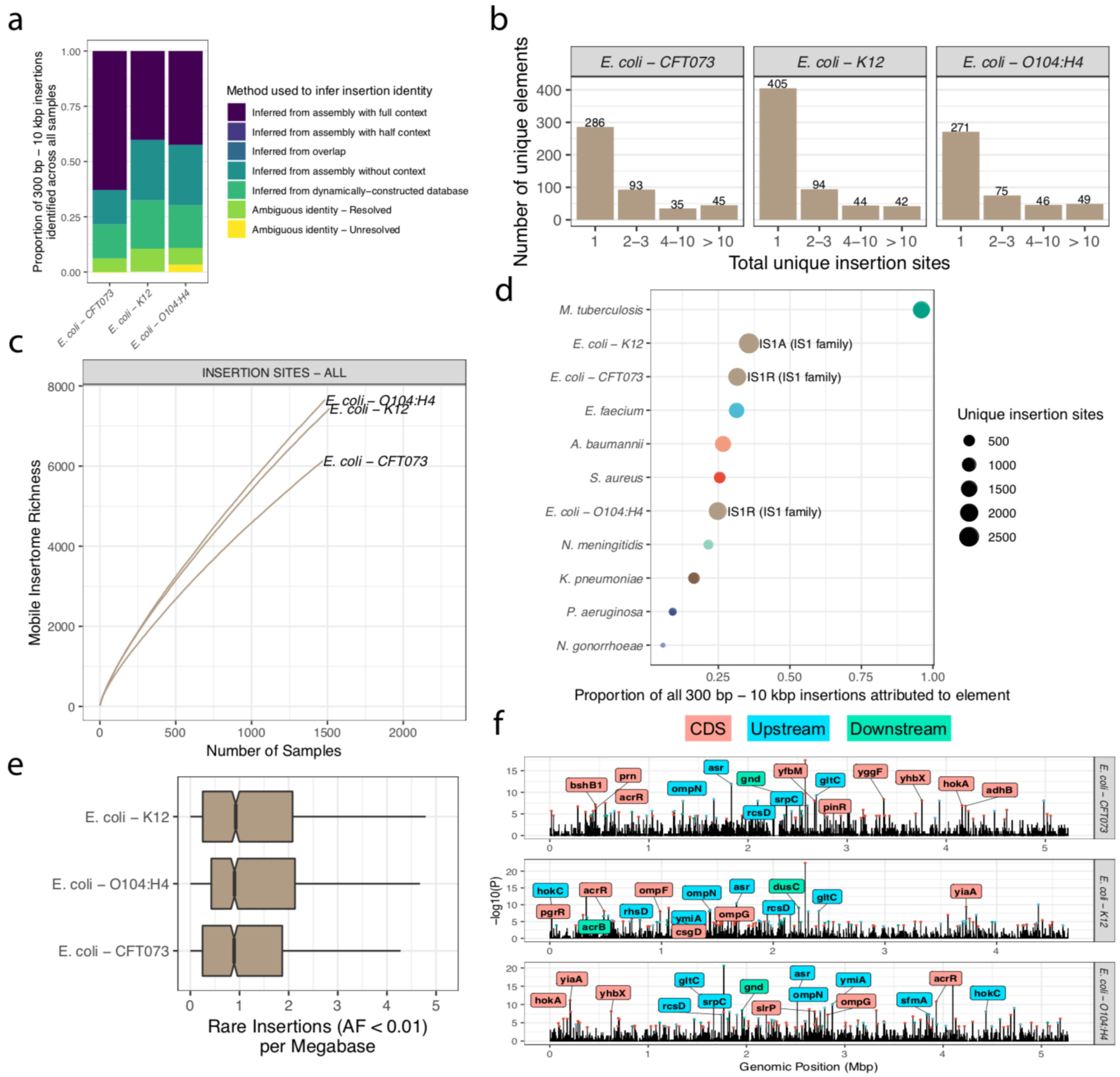


The insertion identification workflow described in this study above was repeated using three different *E. coli* reference genomes: CFT073 (AE014075.1), O104:H4 strain 2011C-3493 (NC_018658.1) and K-12 substrain MG1655 (NC_000913.3) **a**, Results of the workflow described in terms of the types of sequence inference method used to identify each insertion (compare to Figure 1c). **b**, The number of unique sequence elements identified, binned by *E. coli* reference genome and total unique insertion sites (compare to Figure 2a). **c**, The number of new insertions identified as additional isolates are analyzed (compare to Figure 2b). **d**, An analysis of the most MGEs found when using different *E. coli* genomes, defined by the total number of unique insertion sites where the element is found (compare to Figure 2c). **e**, Notched boxplots of the number of rare insertions detected across all samples using each *E. coli* reference genome, adjusted by megabase of genome (compare to Figure 2d). **F**, Analysis of MGE insertion hotspots found when using the three different *E. coli* reference genomes (compare to Figure 4a).

**Supplementary Figure 6: Further demonstration of the sensitivity of the *mustache* insertion identification tool.**

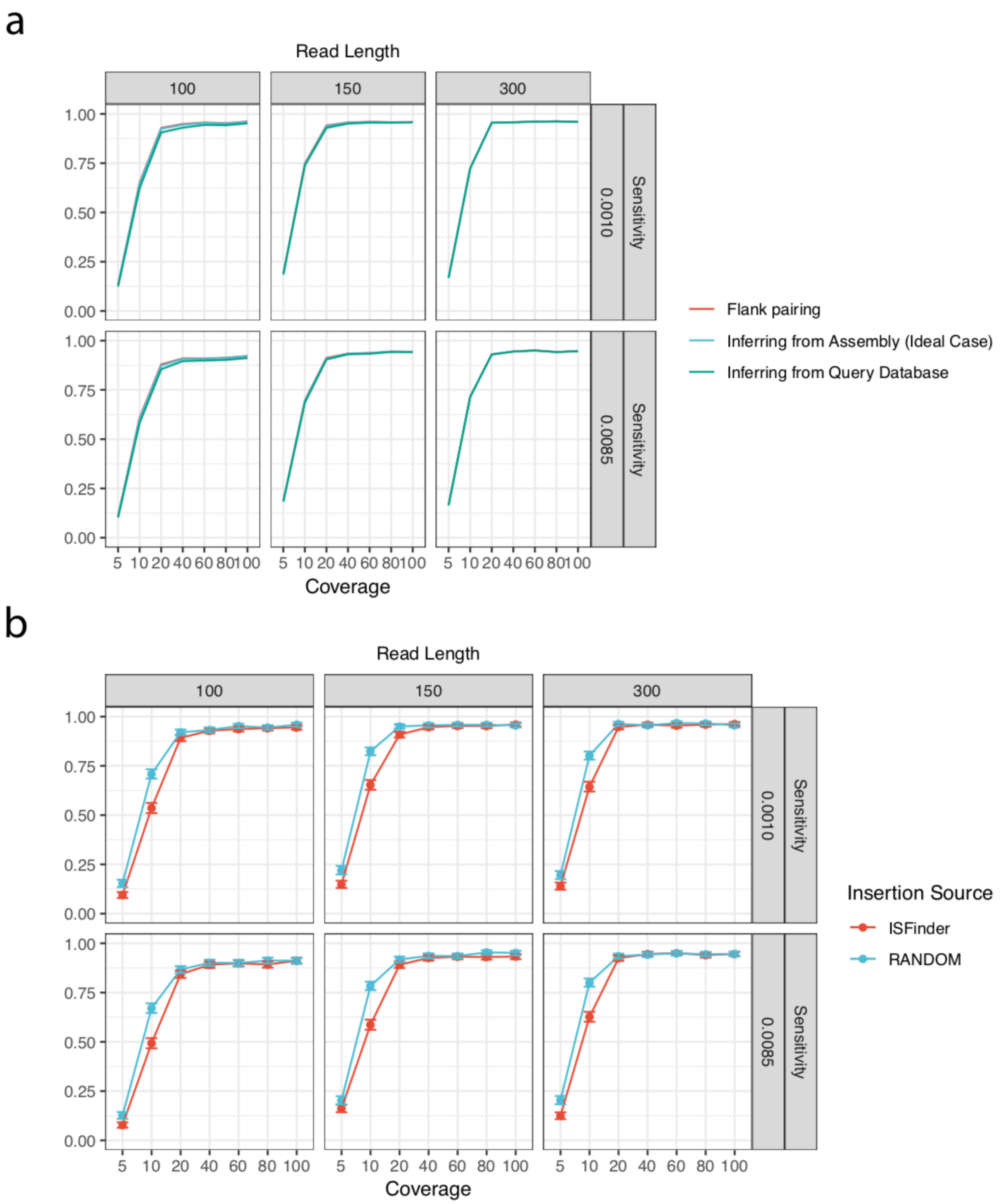


**a**, The sensitivity of different sequence inference steps implemented in *mustache*. "Flank pairing" refers to the sensitivity of mustache prior to any attempt to infer the sequence itself, considered an upper limit on the overall sensitivity. "Inferring from Assembly (Ideal Case)" refers to the scenario where the identity of the inserted sequence is inferred from a perfect, complete assembly of the isolate under study. "Inferring from Query Database" indicates the sensitivity when using a database of known inserted elements with *mustache*. These simulations are not independent of each other, meaning that the exact same simulated genomes were analyzed using the three different sequence inference approaches. For the most part, the curves overlap completely with each other, with the exception of coverages below 40, where inferring from the assembly and inferring from the query database are slightly less sensitive. **B**, Comparing the sensitivity of *mustache* when identifying species-specific elements found in the ISfinder database, and when identifying randomly generated elements (RANDOM). At low coverage, *mustache* is more sensitive to identifying random elements than species-specific elements. This is most likely because random elements do not already exist in the reference genome, and when the inserted element also exists within the reference genome it can pull away reads mapping to the insertion junction. This can be avoided by masking known MGEs from the read aligner being used.

**Supplementary Table 1: Comparison of the features of different previously published MGE insertion identification tools.**

A table giving a brief description of the features offered by different MGE insertion identification tools. Compares the tools *mustache* (the tool presented in this study), *panISa*, *ISEScan*, *breseq*, *ISMapper*, *ITIS*, *ISSeeker*, and *ISQuest*.

**Supplementary Table 2: Samples downloaded randomly from NCBI SRA database for nine pathogens.**

This includes all the run accession, sample accession, experiment excession, and study accession information for all of the samples analyzed for the first part of this study. The species and reference genome are also specified. Also included are the *E. coli* samples that were aligned to the O104:H4 and K-12 reference genomes for comparison with the CFT073 reference genome.

**Supplementary Table 3: Summary table describing the inserted element clusters identified for each species.**

All the inserted element clusters identified in this study are summarized, where each row describes a single cluster. Each column is described as follows: “species” indicates the species where the cluster was identified; “genome” indicates the reference genome used in the *mustache* pipeline; “cluster” is an identifier to describe the specific sequence cluster when all identified insertions are clustered using CD-HIT-EST at 90% sequence identity across 85% of their sequence; “cluster_repr_id” refers to the sequence used as a representative of the cluster; “cluster_repr_seqlength” indicates the length of the cluster's representative sequence; "phmmer_transposase" indicates the type of transposase found in the element, if any, as predicted by using the ISEScan pHMM database (Xie and Tang 2017); "resfinder_genes" indicates the antibiotic resistance genes found by ResFinder within the sequence cluster, with the number in parentheses indicating the number of copies of the gene (Zankari et al. 2012); "prokka_genes" indicates the genes predicted to exist in the sequence cluster, with "tnp" indicated a transposase, and "hypo" indicate a hypothetical protein, and the number of copies of each gene in parentheses; "unique_sites" is the number of unique sites where this element is predicted to be found across all analyzed isolates for the species; "prop_samples" indicates the proportion of isolates from this species where at least one insertion belonging to this cluster was found; "avg_copy" indicates the average number of insertions associated with this cluster when at least one has been found; "prop_IAwFC" indicates the proportion of all insertions of this sequence cluster that were identified according to the "Inferred from assembly with full context" approach; "prop_IAwHC" indicates the proportion of all insertions of this sequence cluster that were identified according to the "Inferred from assembly with half context" approach; "prop_IO" indicates the proportion of all insertions of this sequence cluster that were identified according to the "Inferred from overlap" approach; "prop_IAwoC" indicates the proportion of all insertions of this sequence cluster that were identified according to the "Inferred from assembly without context" approach; "prop_IDB" indicates the proportion of all insertions of this sequence cluster that were identified according to the "Inferred dynamically-constructed database" approach; "prop_AMrASC" indicates the proportion of all insertions of this sequence cluster that were initially ambiguous (See Supplementary Figure 1f), but were resolved by comparison with all other insertions found at that site across all samples; "prop_AMrMS" indicates the proportion of all insertions of this sequence cluster that were initially ambiguous (See Supplementary Figure 1f), but were resolved by prioritizing clusters that were high-confident mobile elements; "prop_AMrML" indicates the proportion of all insertions of this sequence cluster that were initially ambiguous (See Supplementary Figure 1f), but were resolved by prioritizing clusters that were medium-confidence mobile elements; "prop_AM" indicates the proportion of all insertions of this sequence cluster that were ambiguous and could not be resolved; "cluster_repr_seq" is the full inferred sequence of the representative for the cluster.

**Supplementary Table 4: Details of all of the identified insertions across all studied isolates.**

This table includes all of the insertion information for the isolates analyzed in the first part of this study. The columns are described as follows: "species" indicates the species in which the insertion was identified; "genome" indicates the reference genome used in the *mustache* pipeline; "run_accession" is the accession number for the sequence file available for download from the NCBI SRA database; "contig" refers to the name of the reference genome where the insertion was found to exist; "pos_5p" indicates where the soft-clipped reads running from the 5' to 3' direction begin with respect to the reference, an indication of the exact insertion site; "pos_3p" is the same as "pos_3p" but for the clipped reads running in the 3' to 5' direction, with the distance between "pos_3p" and "pos_5p" being the length of the direct repeat created; "cluster" indicates the sequence cluster inferred to be inserted at the position; "AF" is the allelic frequency of this specific insertion across all analyzed isolates for the species; "softclip_count_5p" is the number of reads found to support this insertion running from the 5' to the 3' direction; "softclip_count_3p" is the number of reads found to support this insertion running form the 3' to 5' direction; "total_count_5p" is the total number of reads overlapping the 5' insertion site; "total_count_3p" is the total number of reads overlapping the 3' insertion site; "spanning_count" is the total number of reads that span both the 5' and 3' insertion sites by more than 10 base pairs.

**Supplementary Table 5: A mapping of all of the identified sequences to their cluster assignments.**

Many of the cluster sequences identified contain multiple sequences that are similar, but with slight differences. This likely reflects the natural variation of these inserted elements across isolates. This table maps each unique sequence to its assigned cluster, and the columns are described as follows: “species” indicates the species where the cluster was identified; “genome” indicates the reference genome used in the *mustache* pipeline; “cluster” is an identifier to describe the specific sequence cluster when all identified insertions are clustered using CD-HIT-EST at 90% sequence identity across 85% of their sequence; “cluster_repr_id” refers to the sequence used as a representative of the cluster; “seq_id” is an identifier used to refer to the sequence in the “seq” column; “seq” is the full sequence of this member of the cluster.

**Supplementary Table 6: Predicted ORFs for each reference genome.**

All coding sequences predicted by Prokka and used as genome annotations in this study. Includes species, reference genome, coordinates of coding sequence in BED format, the gene name, and the gene description as assigned by Prokka.

**Supplementary Table 7: Summary of the MGE insertion hotspots identified and represented Figure 4a.**

All the significant MGE insertion hotspots identified and represented in Figure 4a. Each row describes a single MGE insertion hotspot. The columns are described as follows: “species” indicates the species where the hotspot was identified; “genome” indicates the reference genome used in the *mustache* pipeline; “contig” refers to the name of the reference genome where the hotspot was found; “start” is the beginning zero-indexed position of the start of the hotspot; “end” is the one-indexed position of the end of the hotspot; “pvalue” is the p-value of the dynamic one-sided exact Poisson test use to identify significant hotspots; “unique_insertion_count” is the number of unique insertions found within the window across all analyzed isolates for the species of interest; “closest_gene” is the coding sequence that is found to be closest to the center of the insertion hotspot; “closest_gene_start” is the zero-indexed start of the closest gene with respect to the reference genome; “closest_gene_end” is the one-indexed end of the closet gene; "distance_to_closest_gene" is the distance from center of the hotspot to the closest gene, with 0 indicating that the hotspot center directly overlaps the closest gene, negatives number indicating that the hotspot center is upstream of the closest gene, and positive numbers indicating that the hotspot center is downstream of the closest gene; "closest_gene_name" is the name of the closest gene as assigned by the Prokka annotation software; "closest_gene_desc" is a description of the closest gene.

**Supplementary Table 8: GO term enrichment results.**

All of the GO term enrichment results, a subset of which are displayed in Figure 4b. The columns are described as follows: "species" indicates the species in which the hotspots were identified; "genome" indicates the reference genome used in the *mustache* pipeline; "id" refers to the GO term ID being tested; "name" is the description associated with the GO term ID; "namespace" is the GO namespace used for the term, including "cellular_component", "biological_process", and "molecular_function"; "hotspot_goterm" refers to the number of coding sequences in the reference genome that fall within the GO term and are near MGE insertion hotspots; "hotspotal total" refers to the total number of MGE insertion hotspots found in the species' genome; "background_goterm" refers to the total number of coding sequences in the reference genome that were assigned to the GO term; "background total" refers to the total number of sequences used as a background for the hypergeometric test; "hypergeometric_pval" is the p-value of the hypergeometric test used to identify GO terms with coding sequences that are enriched near hotspots; "log2FC" refers to the $\log_2$(Fold Change) enrichment of the GO term; "FDR_qval" refers to the the FDR *q*-value of the test as calculated by the *qvalue* package in R (Dabney, Storey, and Warnes 2010); "hotspot_goterm_genes" indicates all of the coding sequences within the species' reference genome that fall within the GO term and are near hotspots; "goterm_genes" indicates all of the coding sequences within the species' reference genome.

**Supplementary Table 9: HOMER target-sequence motifs for mobile genetic elements.**

A description of the results of the HOMER motif enrichment analysis, using each clusters insertion sites to identify motifs. Each row describes a motif. Each column is described as follows :“species” indicates the species in which the motifs were identified; “genome” indicates the reference genome used in the *mustache* pipeline; “cluster” indicates the sequence cluster that is associated with the discovered motif; “motif_cluster” refers to the cluster of identified motifs to which this motif belongs; “motif_consensus” is a text representation of the identified motif; “log_p_value” is the log(*P*) statistic associated with the motif; “tgt_num” is the number of insertion sites HOMER searched for common motifs; “tgt_pct” refers to the percentage of target sites that contain the identified motif; “bgd_pct” refers to the percentage of randomly chosen background sites that contain the motif; “motif_file” refers to the file location of a sequence logo file in PNG format to represent the target-sequence motif can be found, these locations are with respect to the ZIP file Supplementary File 1.

**Supplementary Table 10: Genes disrupted by IS insertion in Baym et al. Megaplate intermediate-step trimethoprim megaplate experiment.**

A list of all the IS insertions identified in the Baym et al. Megaplate intermediate-step trimethoprim megaplate experiment. Each row is an identified insertion, which includes the sample name, the sample run accession, the position of the insertion, the IS element that is inserted, and the gene that is presumed to be affected by the insertion.

**Supplementary Table 11: Summary of Multidrug resistant (MDR) *E. coli* isolate samples.**

Summary of the samples downloaded from the NCBI SRA database and included as part of the MDR collection of *E. coli* isolates. Includes the SRA accession information, and the antibiotic resistance phenotypes for each isolate. For the purposes of this experiment, “intermediate” was considered grouped together with “resistant” to form a binary variable.

**Supplementary Table 12: Results of GWAS of *E. coli* isolates in MDR and Hospital collections.**

Significant results (FDR $\leq$ 0.05) of the GWAS of large insertions and antibiotic resistance phenotypes for the MDR and Hospital collections. Each row is a single association test. Each column is described as follows: “collection” refers to the collection of isolates analyzed, either “HOSPITAL” or “MDR”; “antibiotic” refers to the antibiotic resistance phenotype tested; “test_type” indicates if the association test was either the presence/absence of a given “insertion”, or the presence/absence of any insertion within a given “region”; “label” is the label used to describe the insertion or region location in terms of gene names; “test” is a descriptor of the specific test performed, including either the insertion information or the details of the region being tested; “p_lrt” is the likelihood-ratio test p-value as calculated by GEMMA; “FDR_qval” refers to the the FDR *q*-value of the test as calculated by the *qvalue* package in R (Dabney, Storey, and Warnes 2010); “beta” is the coefficient associated with the predictor used in the linear-mixed model implemented by GEMMA; “present_case” indicates the number of antibiotic-resistant isolates where the insertion site or region contains an insertion; “present_control” indicates the number of antibiotic-susceptible isolates where the insertion site or region contains an insertion; “absent_case” indicates the number of antibiotic-resistant isolates where the insertion site or region does not contain an insertion; “absent_control” indicates the number of antibiotic-susceptible isolates where the insertion site or region does not contain an insertion; “abs_or” is the |log(odds ratio)| of the test; “N” is the total number of isolates included in the test, where isolates may be missing due to low coverage at the insertion site or due to a lack of antibiotic resistance information.

**Supplementary File 1: All sequence logos for the HOMER target-sequence motifs.**

This includes of all of the significant target-sequence motifs identified by HOMER in PNG format. For information about these motifs, see Supplementary Table 9.
