## Supplementary figures and images for "Mobile genetic element insertions drive antibiotic resistance across pathogens"

### motif1::287--Pseudomonas_aeruginosa::GCF_000006765_1_ASM676v1_genomic::Cluster187::GTCCGCTDCYGG.png

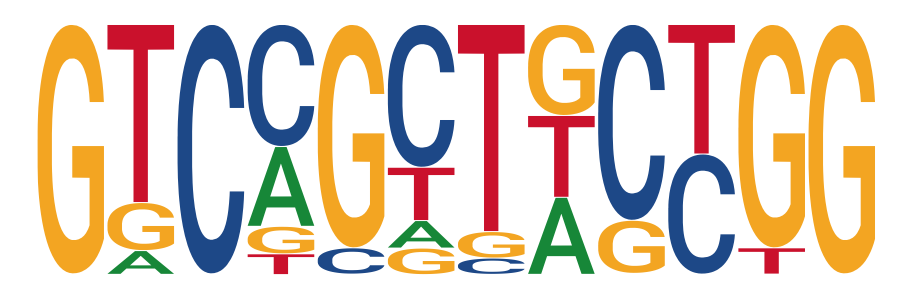

### motif1::287--Pseudomonas_aeruginosa::GCF_000006765_1_ASM676v1_genomic::Cluster188::GGCCGTAAGTGG.png

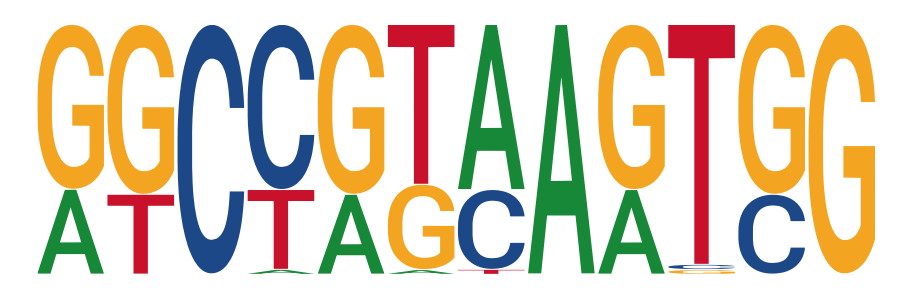

### motif1::287--Pseudomonas_aeruginosa::GCF_000006765_1_ASM676v1_genomic::Cluster211::TTATTCGCCCTA.png

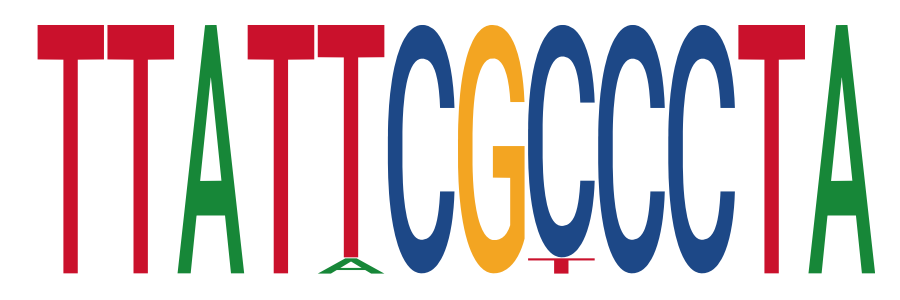

### motif1::470--Acinetobacter_baumannii::GCF_000746645_1_ASM74664v1_genomic::Cluster73::GCGGTAGGGCGA.png

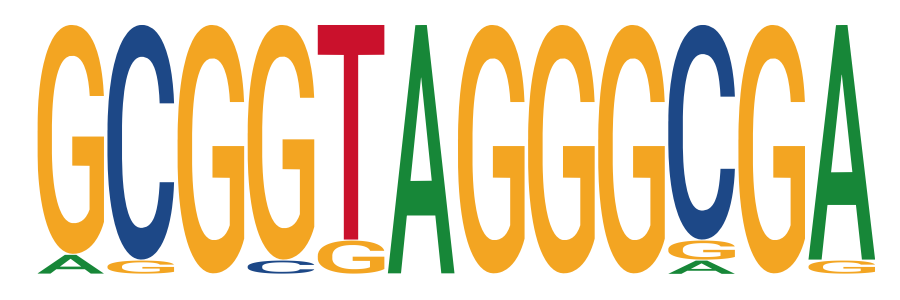

### motif1::470--Acinetobacter_baumannii::GCF_000746645_1_ASM74664v1_genomic::Cluster109::AGTCTTAGTTGA.png

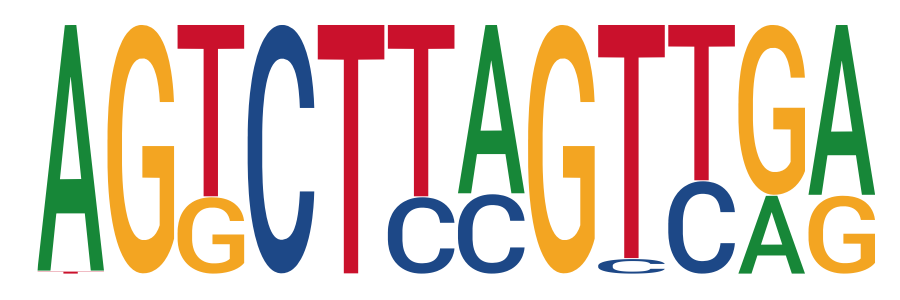

### motif1::470--Acinetobacter_baumannii::GCF_000746645_1_ASM74664v1_genomic::Cluster133::ATAATATATA.png

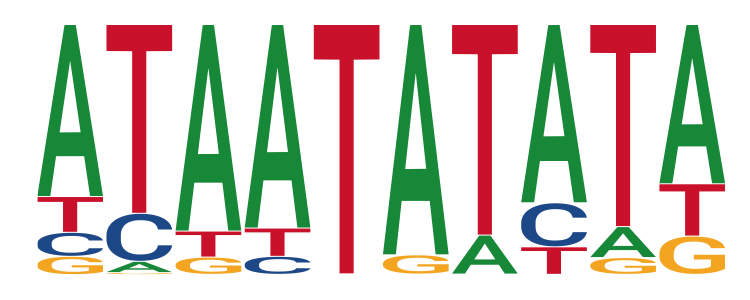

### motif1::470--Acinetobacter_baumannii::GCF_000746645_1_ASM74664v1_genomic::Cluster135::ACTAWTTA.png

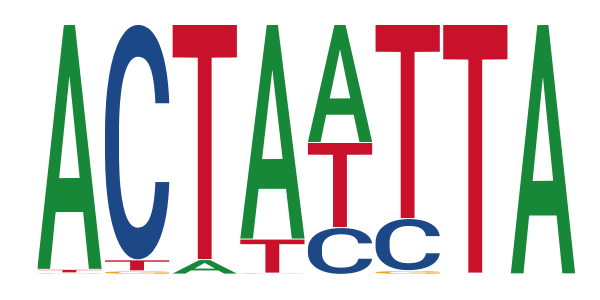

### motif1::487--Neisseria_meningitidis::GCF_000008805_1_ASM880v1_genomic::Cluster31::ATTCCCGCST.png

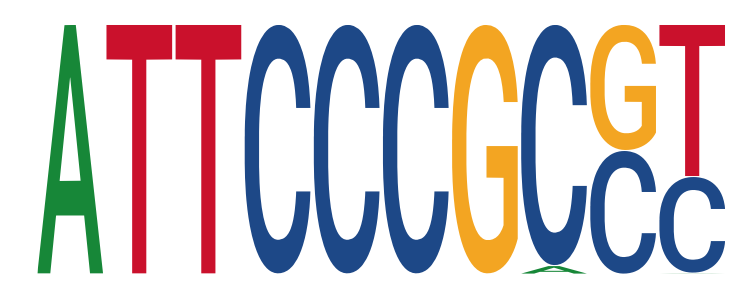

### motif1::487--Neisseria_meningitidis::GCF_000008805_1_ASM880v1_genomic::Cluster364::AAATATTTTATT.png

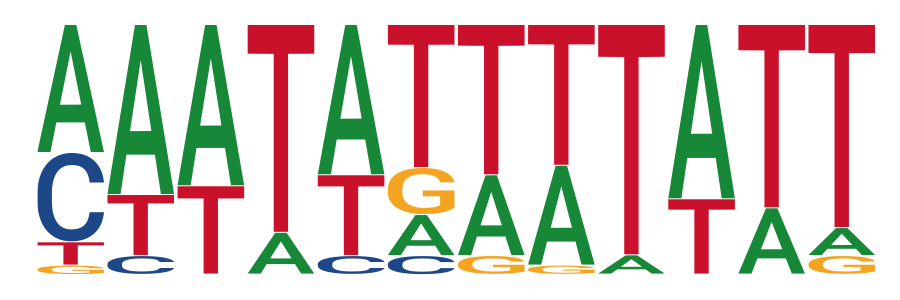

### motif1::562--Escherichia_coli::CFT073::Cluster234::TATATACAGTAT.png

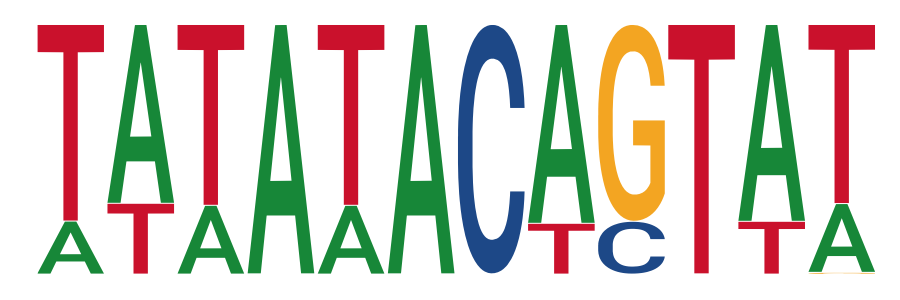

### motif1::562--Escherichia_coli::CFT073::Cluster256::CGCCTTATCCGG.png

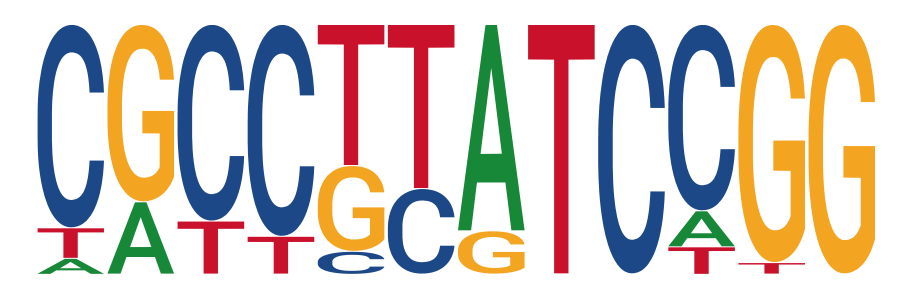

### motif1::573--Klebsiella_pneumoniae::GCF_000240185_1_ASM24018v2_genomic::Cluster189::AGCGCCGCCGGA.png

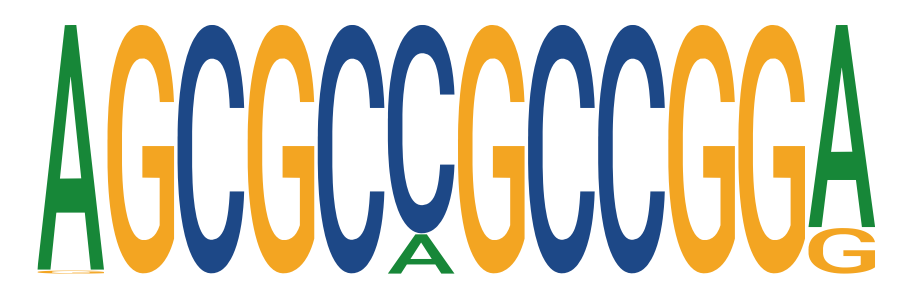

### motif1::573--Klebsiella_pneumoniae::GCF_000240185_1_ASM24018v2_genomic::Cluster234::TNNGCTAABDCW.png

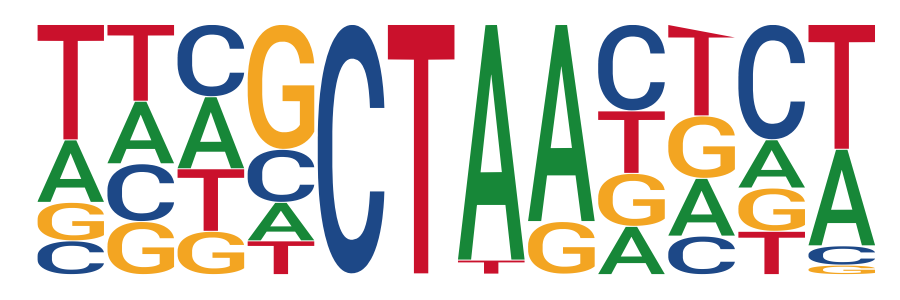

### motif1::573--Klebsiella_pneumoniae::GCF_000240185_1_ASM24018v2_genomic::Cluster240::CTCCTGWTRTTG.png

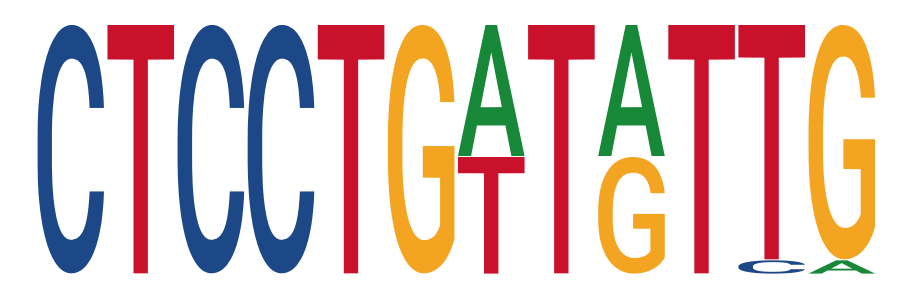

### motif1::573--Klebsiella_pneumoniae::GCF_000240185_1_ASM24018v2_genomic::Cluster247::TTTACATAGCTC.png

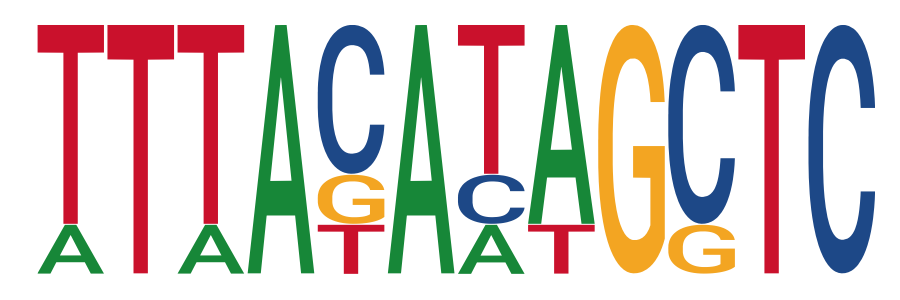

### motif1::1352--Enterococcus_faecium::GCF_000174395_2_ASM17439v2_genomic::Cluster38::TCTTTTWTCGAT.png

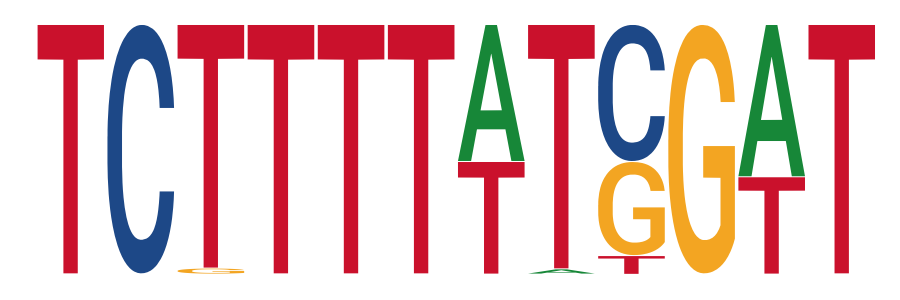

### motif1::1352--Enterococcus_faecium::GCF_000174395_2_ASM17439v2_genomic::Cluster39::TTTAAA.png

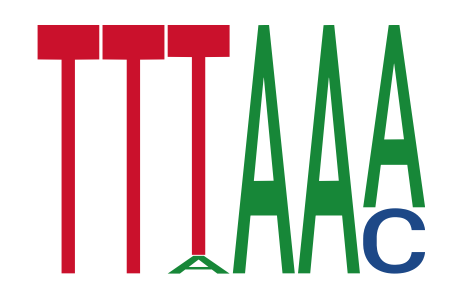

### motif1::1352--Enterococcus_faecium::GCF_000174395_2_ASM17439v2_genomic::Cluster44::TTTAGGAGGA.png

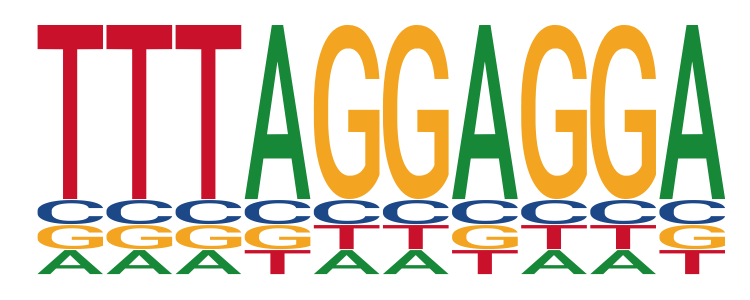

### motif1::1352--Enterococcus_faecium::GCF_000174395_2_ASM17439v2_genomic::Cluster52::GAGAGG.png

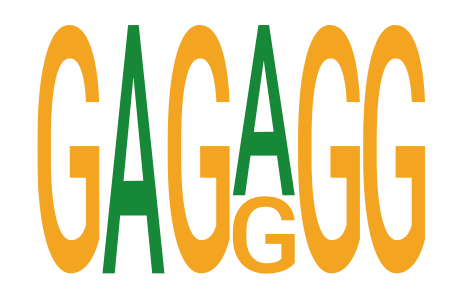

### motif1::1352--Enterococcus_faecium::GCF_000174395_2_ASM17439v2_genomic::Cluster54::GTCACAGCCC.png

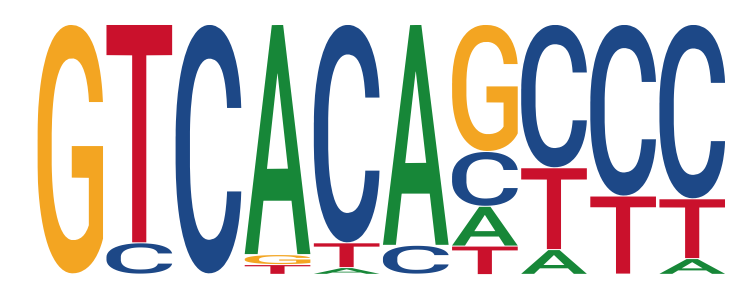

### motif1::1773--Mycobacterium_tuberculosis::GCF_000195955_2_ASM19595v2_genomic::Cluster31::TTTCCGTCCC.png

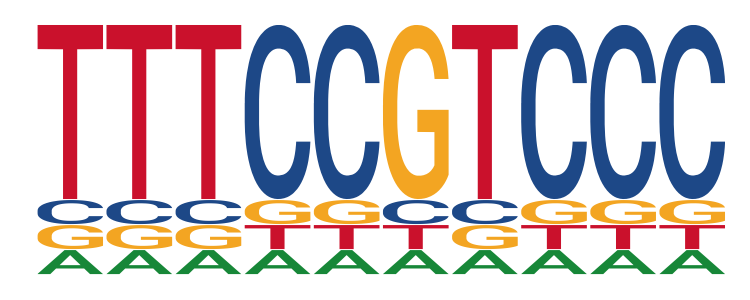

### motif2::487--Neisseria_meningitidis::GCF_000008805_1_ASM880v1_genomic::Cluster319::TTTCAAATGSCS.png

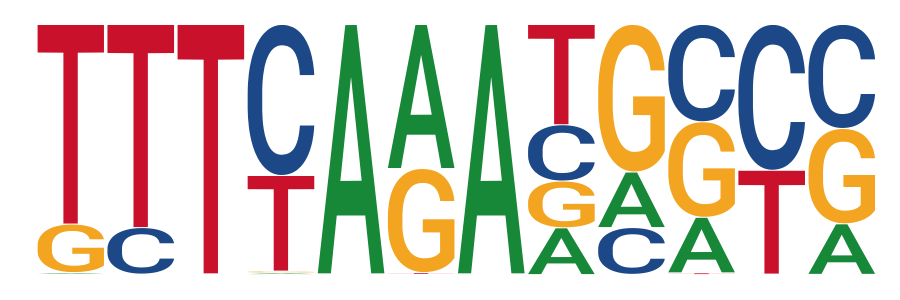

### motif2::562--Escherichia_coli::CFT073::Cluster298::RCTNAGCRTBYA.png

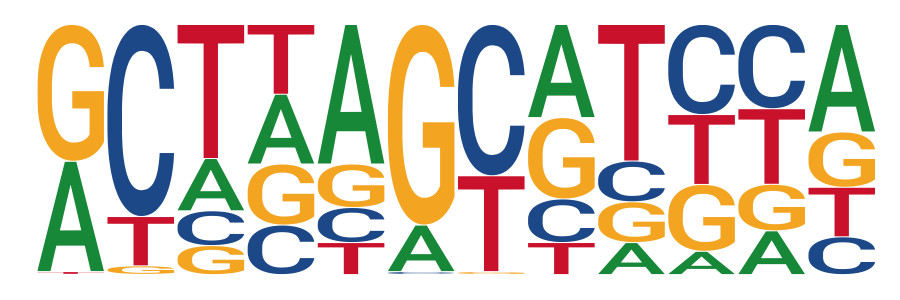

### motif2::562--Escherichia_coli::CFT073::Cluster390::GSSBTTTTTTTT.png

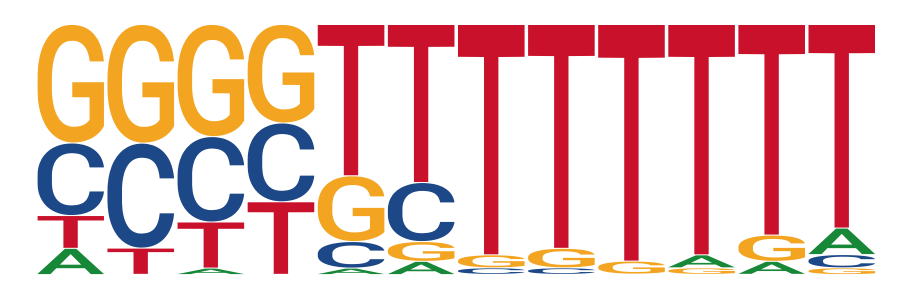

### motif2::1352--Enterococcus_faecium::GCF_000174395_2_ASM17439v2_genomic::Cluster53::GTCCCAGCCTCT.png

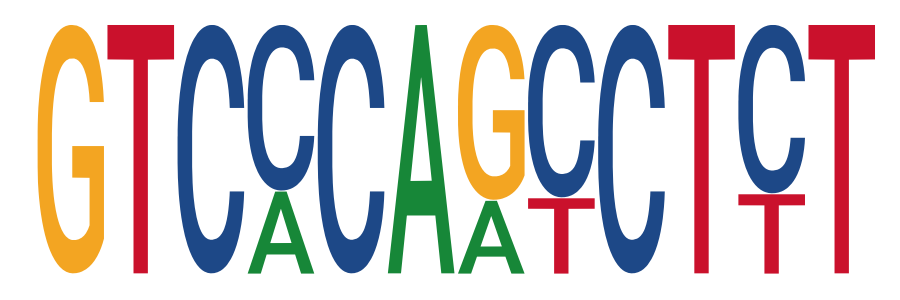

### motif3::487--Neisseria_meningitidis::GCF_000008805_1_ASM880v1_genomic::Cluster339::CTAGTM.png

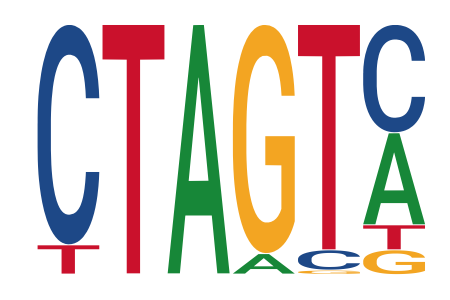

### motif3::562--Escherichia_coli::CFT073::Cluster269::TCGGGCCTACRT.png

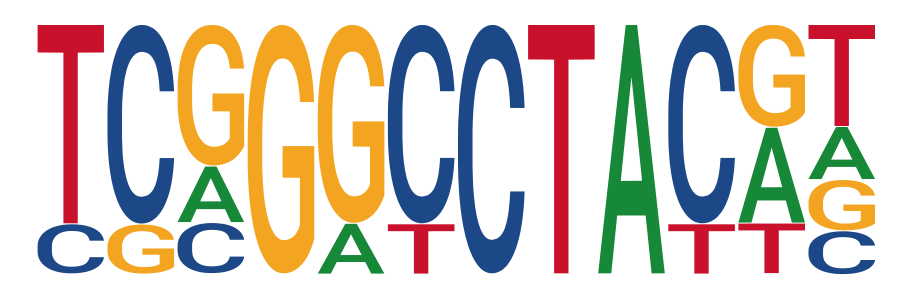

### motif3::573--Klebsiella_pneumoniae::GCF_000240185_1_ASM24018v2_genomic::Cluster213::TATACAGTATAG.png

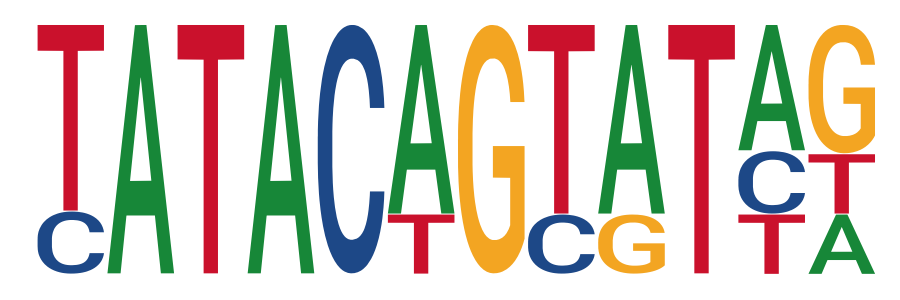

### motif3::1352--Enterococcus_faecium::GCF_000174395_2_ASM17439v2_genomic::Cluster61::GGACTGTGACAA.png

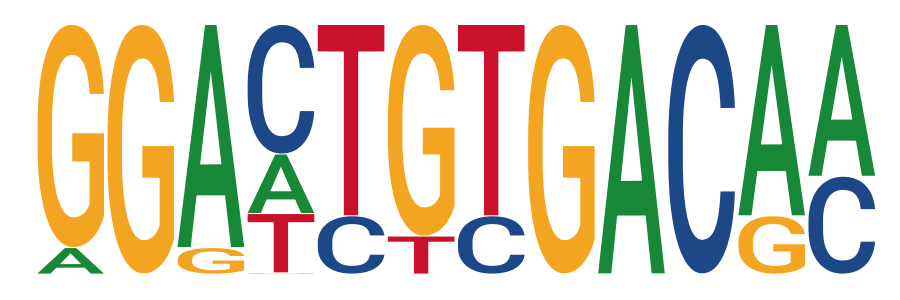

### motif3::1352--Enterococcus_faecium::GCF_000174395_2_ASM17439v2_genomic::Cluster106::TTAA.png

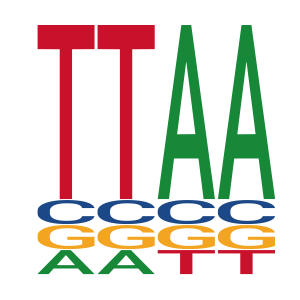

### motif4::470--Acinetobacter_baumannii::GCF_000746645_1_ASM74664v1_genomic::Cluster75::TTATAAAA.png

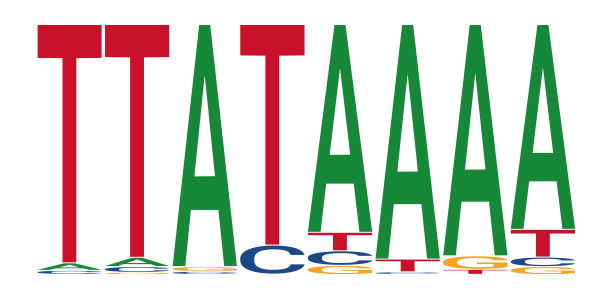

### motif4::573--Klebsiella_pneumoniae::GCF_000240185_1_ASM24018v2_genomic::Cluster208::ATGCTBAGCR.png

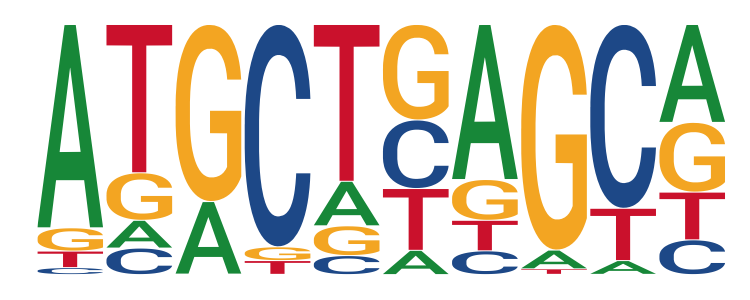

### motif4::1280--Staphylococcus_aureus::GCF_000013425_1_ASM1342v1_genomic::Cluster119::TGTCCCAACCTC.png

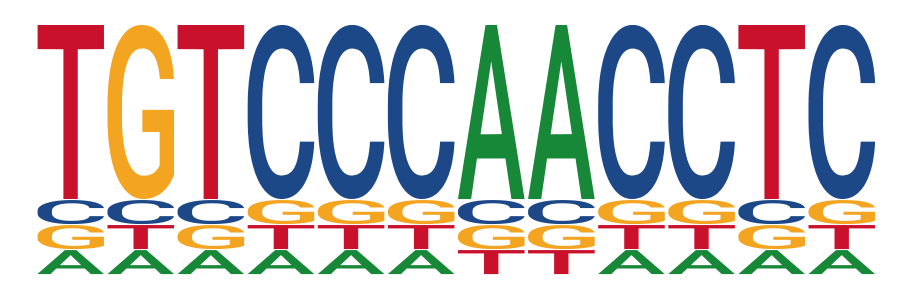

### motif5::470--Acinetobacter_baumannii::GCF_000746645_1_ASM74664v1_genomic::Cluster78::ATGCTSAGCATG.png

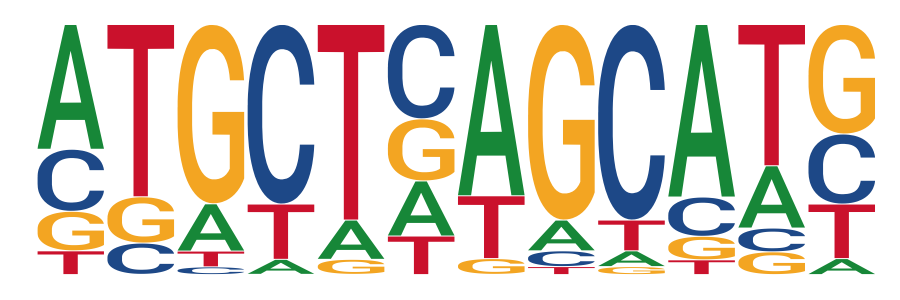

### motif5::562--Escherichia_coli::CFT073::Cluster274::GGTGCCVTTTTT.png

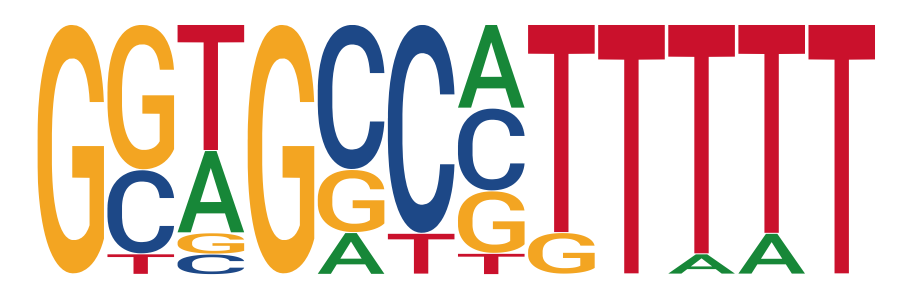

### motif5::562--Escherichia_coli::CFT073::Cluster329::AYWHHNCTAA.png

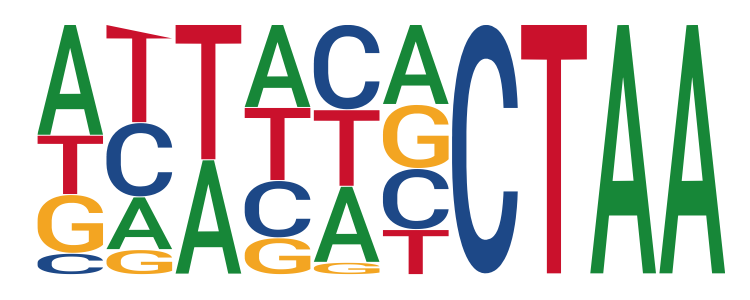

### motif6::573--Klebsiella_pneumoniae::GCF_000240185_1_ASM24018v2_genomic::Cluster241::CTAG.png

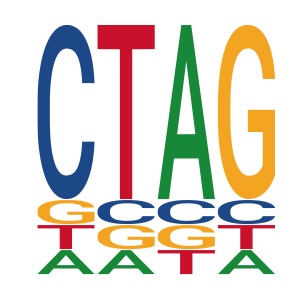

### motif6::1280--Staphylococcus_aureus::GCF_000013425_1_ASM1342v1_genomic::Cluster143::TTTATATAAAAA.png

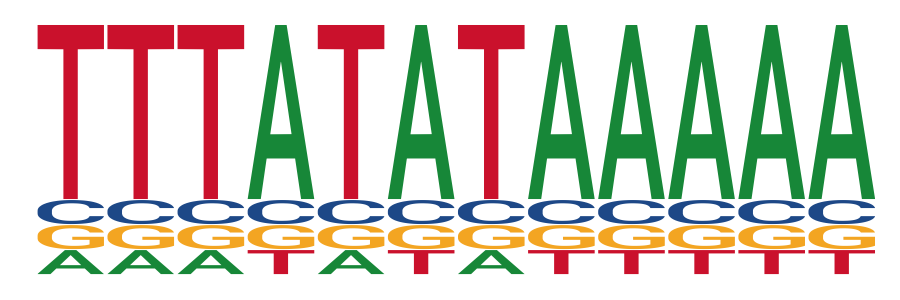

### motif7::287--Pseudomonas_aeruginosa::GCF_000006765_1_ASM676v1_genomic::Cluster199::YCCTAG.png

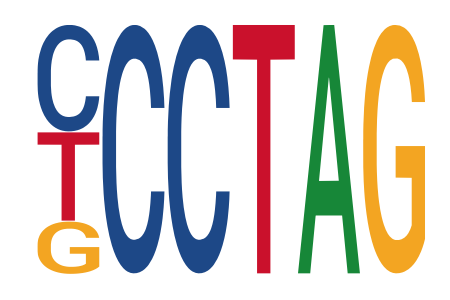

### motif7::287--Pseudomonas_aeruginosa::GCF_000006765_1_ASM676v1_genomic::Cluster266::GBTTTTATTC.png

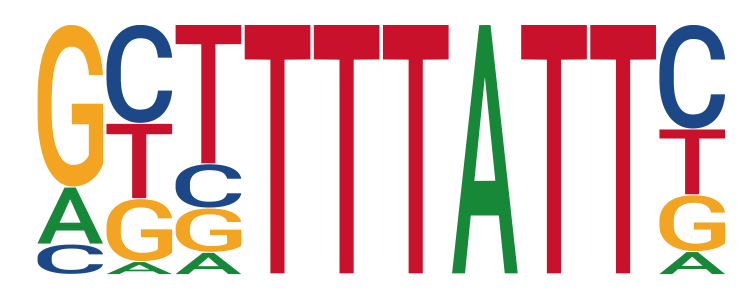

### motif8::287--Pseudomonas_aeruginosa::GCF_000006765_1_ASM676v1_genomic::Cluster258::GASVRCTAGH.png

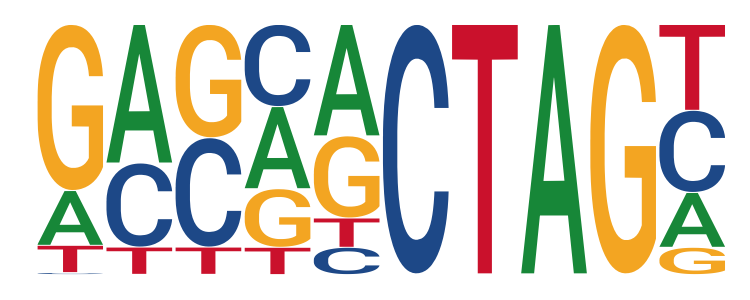

### motif9::287--Pseudomonas_aeruginosa::GCF_000006765_1_ASM676v1_genomic::Cluster216::TRRTGGCTAGKG.png

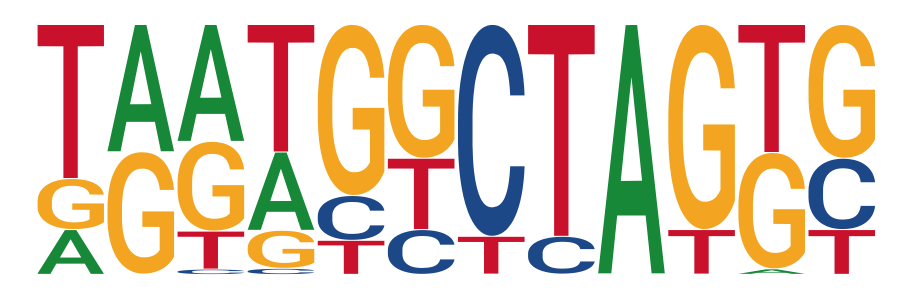

### motif9::1280--Staphylococcus_aureus::GCF_000013425_1_ASM1342v1_genomic::Cluster120::TCTTTTTTTATT.png

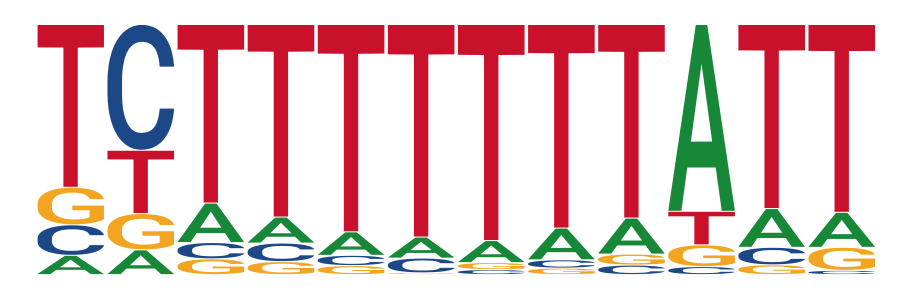

### motif10::1352--Enterococcus_faecium::GCF_000174395_2_ASM17439v2_genomic::Cluster87::TTATAATAAAAA.png

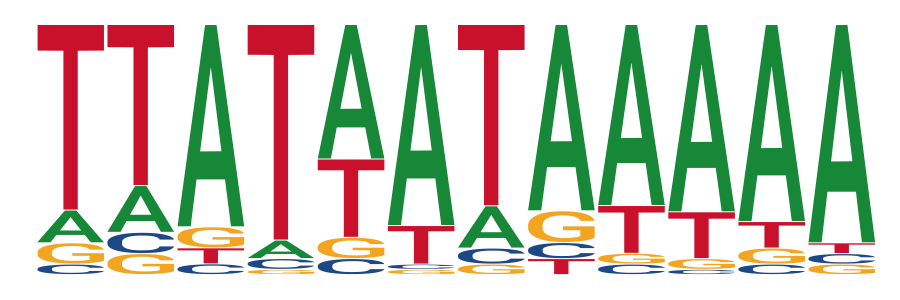

### motif11::1280--Staphylococcus_aureus::GCF_000013425_1_ASM1342v1_genomic::Cluster100::GTTTTTGT.png

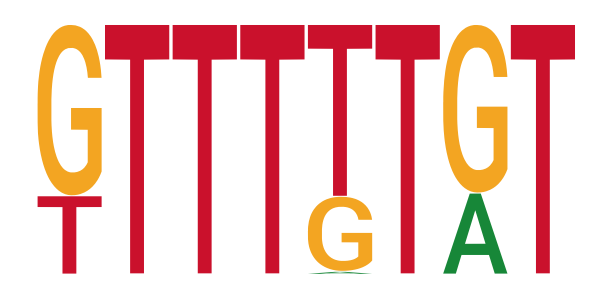

### motif12::470--Acinetobacter_baumannii::GCF_000746645_1_ASM74664v1_genomic::Cluster72::AAAAAACCGCGC.png

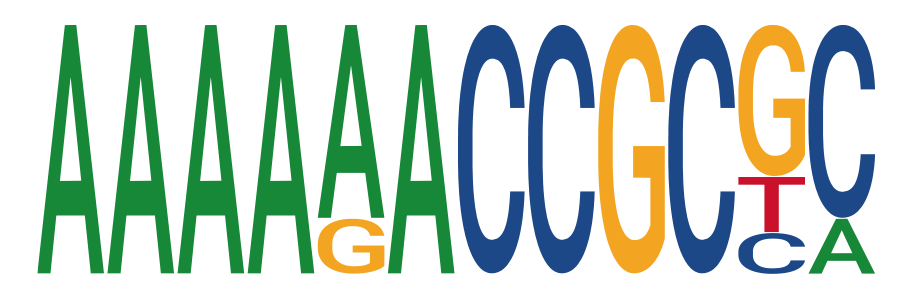

### motif14::1352--Enterococcus_faecium::GCF_000174395_2_ASM17439v2_genomic::Cluster89::TTATATAATAAA.png

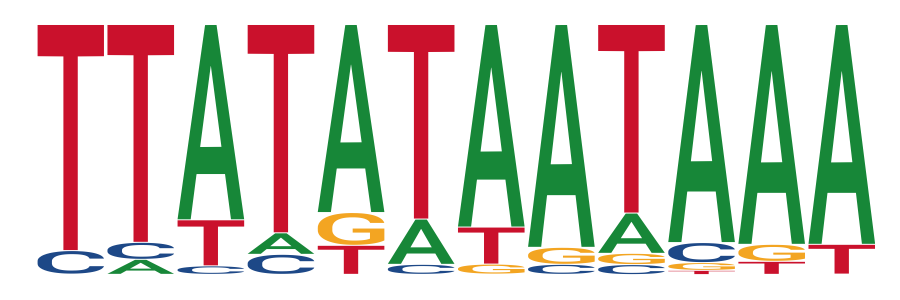

### motif15::562--Escherichia_coli::CFT073::Cluster323::CATGGCGATTTT.png

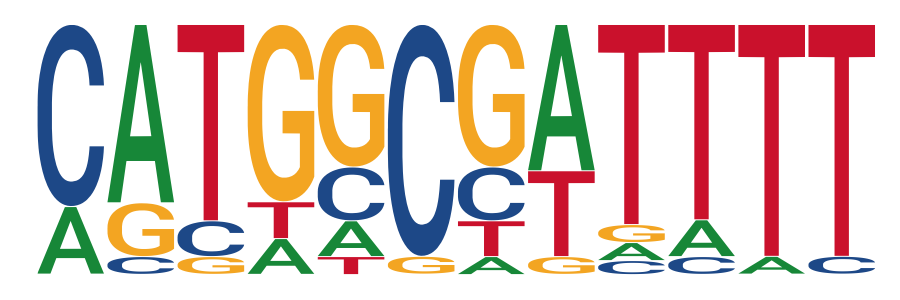

### motif16::287--Pseudomonas_aeruginosa::GCF_000006765_1_ASM676v1_genomic::Cluster222::CGGGYTTTTTTT.png

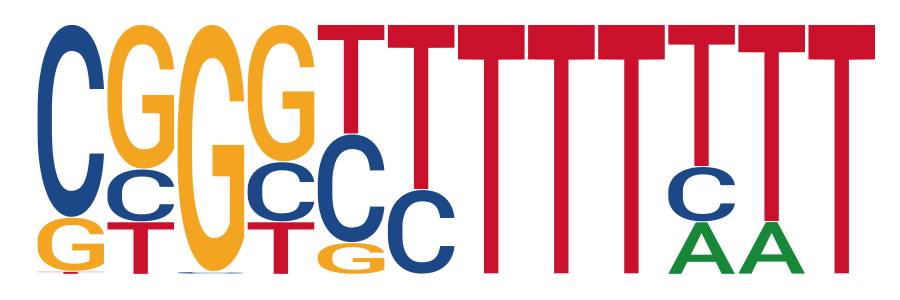

### motif17::1352--Enterococcus_faecium::GCF_000174395_2_ASM17439v2_genomic::Cluster88::TATATAAA.png

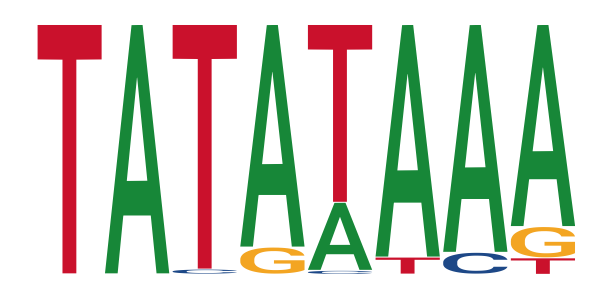

### motif19::1280--Staphylococcus_aureus::GCF_000013425_1_ASM1342v1_genomic::Cluster13::TTTTATWATAAA.png

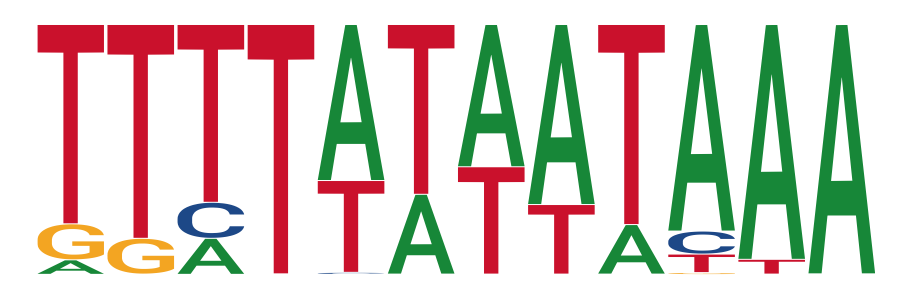

### motif20::1352--Enterococcus_faecium::GCF_000174395_2_ASM17439v2_genomic::Cluster58::AAAAAAAAAA.png

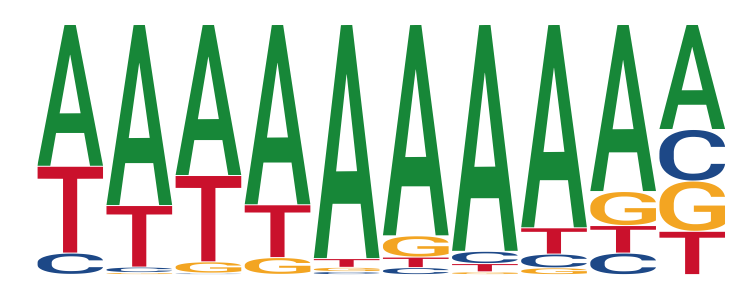

### motif21::287--Pseudomonas_aeruginosa::GCF_000006765_1_ASM676v1_genomic::Cluster223::CGGCCTTTTTTT.png

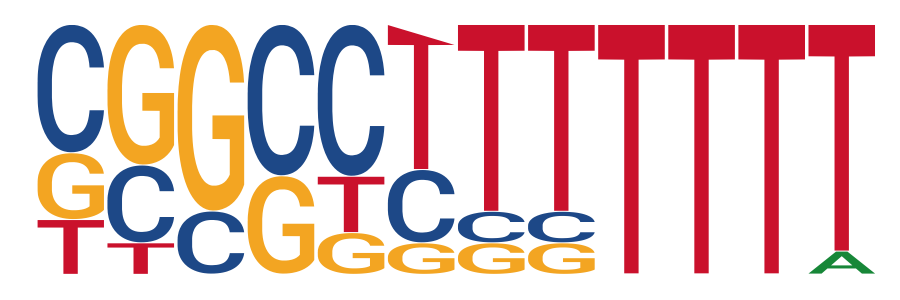

### motif22::470--Acinetobacter_baumannii::GCF_000746645_1_ASM74664v1_genomic::Cluster91::CTCTTTTTTT.png

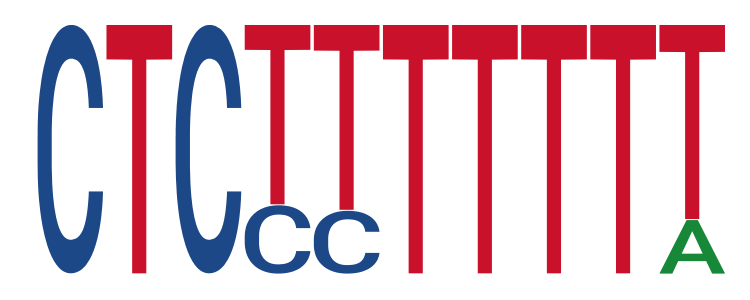

### motif23::562--Escherichia_coli::CFT073::Cluster372::ACGCGATTTT.png

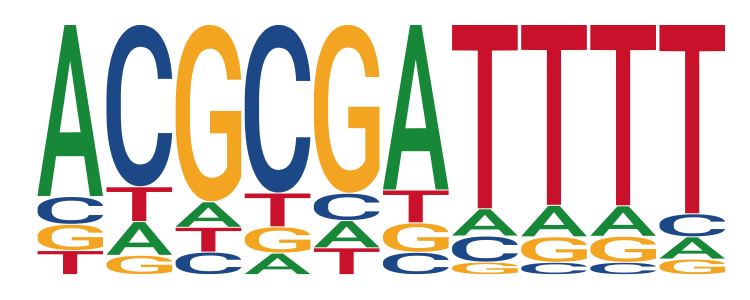
